# Expanding and Decoding the Chemistry of Phospholipid Headgroup in Eukaryotes

**DOI:** 10.64898/2026.01.06.697852

**Authors:** Riku Nakanishi, Airi Matsushita, Yoko Uchino, Yuki Oba, Nanaho Amano, Yuki Kageyama, Kotaro Hirano, Akira Murakami, Nobuhiko Tachibana, Takuma Kishimoto, Sho Nagase, Yoshihiro Ujihara, Kazuki Sugo, Taiki Shigematsu, Tomoki Sato, Shinji Miura, Makoto Inai, Yuji Hara, Masaki Tsuchiya

## Abstract

Cellular membranes have diverse phospholipids, chemical differences in whose headgroups impact many biological processes. Phosphatidylcholine is an essential phospholipid for human health, but not universally required for life. The evolutionary mechanisms underlying phospholipid preferences remain poorly understood, due to the difficulty of investigating metabolite structure–activity relationships in a cellular context. Here, we developed a generalizable metabolic-rewiring method to manipulate phospholipid headgroups together and their biological effects. This approach utilizes synthetic media to hijack evolutionarily conserved phosphatidylcholine biosynthesis, leveraging xenobiotics as principal precursors for scalable headgroup transformations. By identifying over 100 artificial headgroups, we expanded the chemical diversity of xenobiotic phospholipids. Unexpectedly, we discovered that subtle headgroup alterations produced distinct mammalian cellular activities. We demonstrated that chemical headgroup modifications differentially elicited structure-dependent effects on phospholipid–protein interactions, calcium dynamics, transcriptomic profiles, and stem cell differentiation. Notably, cross-species comparison revealed that human and yeast cells have different headgroup preferences critical for cell life and death. As proof-of-concept, interactome analysis identified headgroup-sensitive human microproteins vital for mitochondrial respiration, but non-conserved in yeast. These results exemplify the evolutionary diversity of key phospholipid–protein interactions, illustrating why humans depend on phosphatidylcholine. Overall, our findings establish a programmable platform for elucidating and engineering phospholipid-driven cellular functions.

## Introduction

Membrane lipids are essential for numerous cellular processes^1^. Although a single lipid species can form membrane bilayers that serve as simple physical barriers, cells have structurally diverse phospholipids distinguished by their headgroups^2^ (e.g., phosphatidylcholine [PC] and phosphatidylethanolamine [PE]) (Fig. 1a). The chemical diversity of these headgroups determines membrane physicochemical properties and biological functions, including membrane curvature, surface charge, and lipid–protein interactions^3–5^. Membrane lipid composition also varies widely among organisms^1,3,6^: PC constitutes roughly half of the total lipids in mammals^7^, whereas many bacteria and some yeast strains lack PC entirely^8,9^. Despite PC not being universally required for life, human mutations in PC-biosynthetic enzymes cause diseases that cannot be compensated for by other lipids^2,7,10^. Likewise, abnormal headgroup-forming reactions in other phospholipids lead to various pathologies^2,7,10^. These facts raise fundamental questions about how and why differences in phospholipid headgroups give rise to diverse biological outcomes^2,6,8,11^. A key strategy to answer them is to directly manipulate phospholipid structures within a cellular context, thereby revealing their biological impacts and consequences^12^.

**Figure 1.**
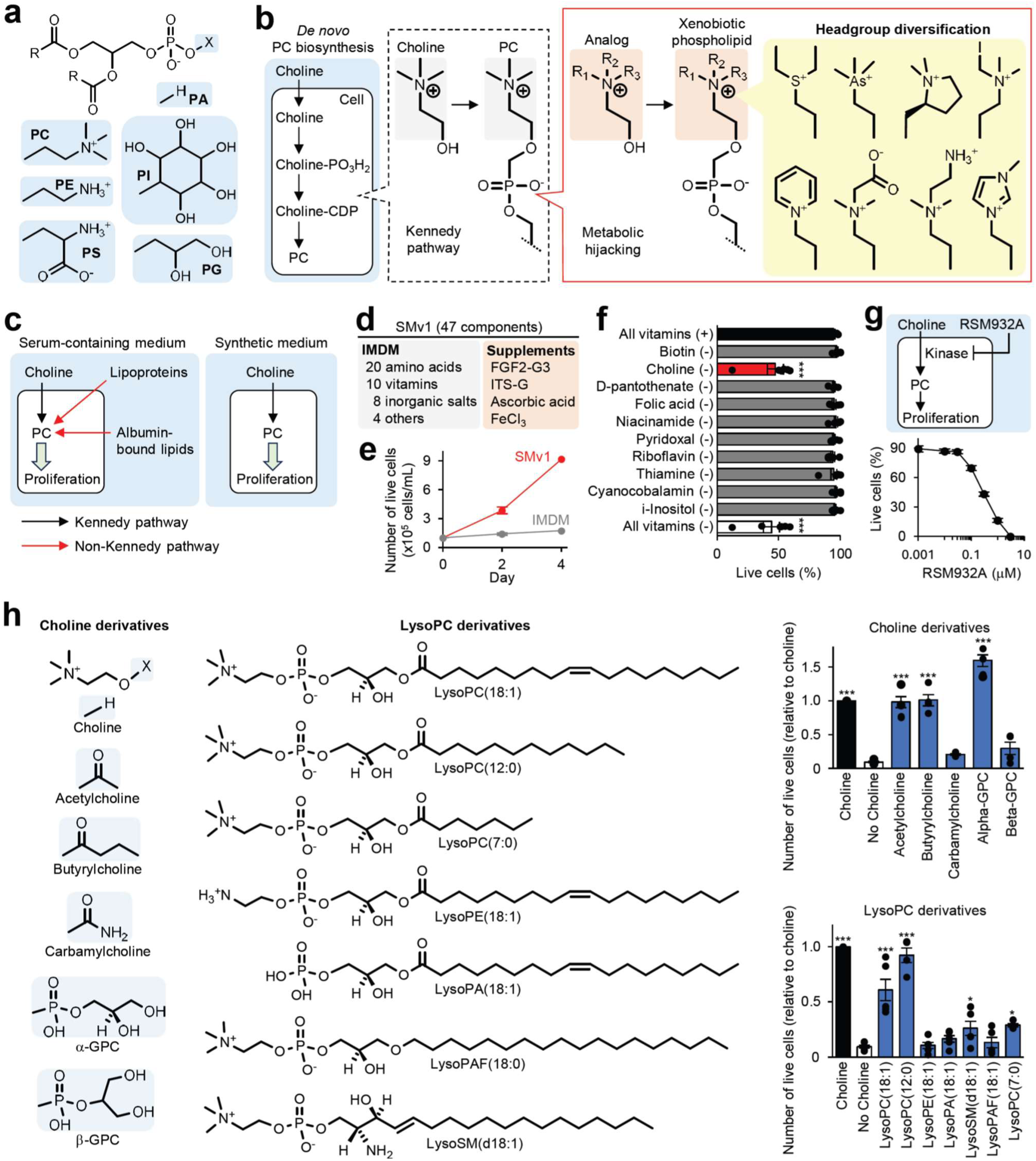
Synthetic media for rewiring phospholipid biosynthetic pathways. (**a**) Headgroups of major glycerophospholipids. PC, phosphatidylcholine; PE, phosphatidylethanolamine; PS, phosphatidylserine; PA, phosphatidic acid; PI, phosphatidylinositol; PG, phosphatidylglycerol. (**b**) Strategy for manipulating phospholipid headgroups. Extracellular choline is transported into cells and converted into the PC headgroup via the Kennedy pathway. Dominating this pathway with choline analogs enables efficient *de novo* biosynthesis of xenobiotic phospholipids with chemically diverse headgroups. (**c**) Serum-containing media supply PC to cells via multiple pathways, whereas synthetic media limit a PC-supplying route to the Kennedy pathway. (**d**) SMv1 consists of IMDM and indicated supplements (detailed composition in Excel file). (**e**) Cell proliferation in SMv1 culture. K562 cells cultured in SMv1 and IMDM were subjected to automated cell counting (n = 6 biological replicates). (**f**) Suppressed proliferation by choline depletion. K562 cells cultured in vitamin-depleted SMv1 for two days were analyzed by cell counting (n = 5 or 6 biological replicates). (**g**) Suppressed proliferation by choline kinase inhibition. K562 cells cultured with RSM932A in SMv1 were analyzed by cell counting (n = 3 biological replicates). (**h**) Suppressed proliferation by choline depletion can be recovered by choline moiety-containing compounds. Structures of choline and lysoPC derivatives are shown. K562 cells cultured with choline derivatives (30 μM) and lysolipid-albumin complexes (10 μM) in choline-depleted SMv1 were analyzed by cell counting (n = 4–6 biological replicates). Data are mean ± SEM. \**p* <0.05, \*\**p* <0.01, \*\*\**p* <0.001 (one-way ANOVA).

However, unlike proteins, lipids are not directly encoded by DNA, and their biosynthesis involves multiple enzymatic steps, making it challenging to generate specific metabolite structures through conventional genetic approaches^12^. Existing methods to introduce defined phospholipids into living cells remain limited, particularly for mammalian cell culture systems in a scalable manner. Direct phospholipid transfer from culture media to cell membranes is generally inefficient due to the poor water solubility. Cyclodextrin can manipulate individual phospholipid species, but it is limited to cell surface^13^. Alternatively, phospholipase D can replace the PC headgroup with alcohol substrates, but the resulting products typically represent a few percent of cellular lipids^14^.

Previously, we developed chemical labeling techniques to trace PC metabolism via the evolutionarily conserved Kennedy pathway^15,16^, which converts choline into the PC headgroup^7^ (Fig. 1b, left and middle panels). Building on this concept, we envisioned that redirecting this pathway toward choline-like xenobiotics could drive large-scale biosynthesis of corresponding headgroups (Fig. 1b, red frame). Furthermore, leveraging a library of such xenobiotics would enable systematic exploration of diverse headgroup chemistries and their biological effects (Fig. 1b, yellow highlighted).

Here, we report a method to manipulate phospholipid headgroups in eukaryotic cells. This employs synthetic media to reprogram cellular metabolism, using xenobiotics as dominant precursors for the Kennedy pathway. We identified over 100 compounds transformable into phospholipid headgroups, and revealed that subtle headgroup modifications elicited distinct effects on mammalian cell proliferation. Our *in vitro* and *in silico* analyses collectively suggest that xenobiotic phospholipids can serve as membrane lipids with altered bilayer properties and biomolecular interactions. Indeed, we found headgroup structure-dependent differential effects on phospholipid remodeling, Ca^2+^ dynamics, and signaling responses. Moreover, phospholipid–protein interactome analysis identified headgroup-sensitive human membrane proteins. Finally, we demonstrated the applicability of this method to rodent primary cardiomyocytes and muscle stem cells, and cross-species comparison further revealed that human and yeast cells have different headgroup preferences critical for cell viability.

## Results

### Optimizing the Kennedy pathway for efficient xenobiotic phospholipid biosynthesis

To enable efficient conversion of xenobiotics into phospholipid headgroups in mammalian cells (Fig. 1b), we sought to establish a culture platform where the Kennedy pathway and other compensatory pathways could be controlled. However, conventional serum-containing media hinder precise manipulation of cellular lipid metabolism due to abundant protein-bound PC and its metabolites, which are difficult to deplete selectively^17–19^ (Fig. 1c, left). To overcome this limitation, we developed a synthetic medium (SMv1) that lacks any lipidic components and thereby enforces phospholipid biosynthesis from exogenously supplied headgroup precursors (Fig. 1c, right). SMv1 was formulated from previous medium compositions^16,20^ and supported proliferation of human K562 cells for four days (Fig. 1d, 1e). To confirm the significance of choline, we performed single-vitamin depletion assays. Among the ten vitamins, only choline removal suppressed proliferation and induced cell death (Fig. 1f). Consistently, choline kinase inhibition produced the same phenotype (Fig. 1g), establishing that PC biosynthesis from extracellular choline is essential for cell viability in SMv1 culture.

We examined whether choline moiety-containing compounds could rescue the suppressed proliferation under choline depletion (Fig. 1h). Acetylcholine and butyrylcholine – both enzymatically hydrolyzed to choline^7^ – restored proliferation to levels comparable to the choline-supplemented control, but esterase-resistant carbamylcholine did not. L-α-Glycerylphosphorylcholine (α-GPC), another choline-releasing compound^21^, also promoted proliferation, while its isomer β-GPC did not. We further examined monoacyl phospholipids. LysoPC(18:1) and its medium-chain analog lysoPC(12:0) showed proliferative effects, consistent with their salvage into PC via non-Kennedy pathways^7,19^ (Fig. 1c, left). However, other lysolipids such as lysoPE(18:1) had no or less rescue effect. These results demonstrate that synthetic media enable metabolic control of the Kennedy pathway, validating the SMv1 culture system as a promising platform for xenobiotic phospholipid biosynthesis.

### Xenobiotic phospholipids can serve as an alternative to PC

To systematically explore xenobiotic precursors for phospholipid headgroups and their biological effects, we treated K562 cells with choline analogs (30 μM) in SMv1 for lipid characterization and structure–activity relationship analysis (Fig. 2a). The resulting xenobiotic phospholipids were detected by thin-layer chromatography (TLC) and mass spectrometry (MS) (Fig. 2b, in yellow; S1a, S1b). Proliferative activities of the tested analogs were quantified by cell counts normalized to the choline-supplemented control; analogs with proliferation scores >0.3 were defined as active in this setting (Fig. 2b, in red), which exceeded that of the choline-depleted control (score: ∼0.1).

**Figure 2.**
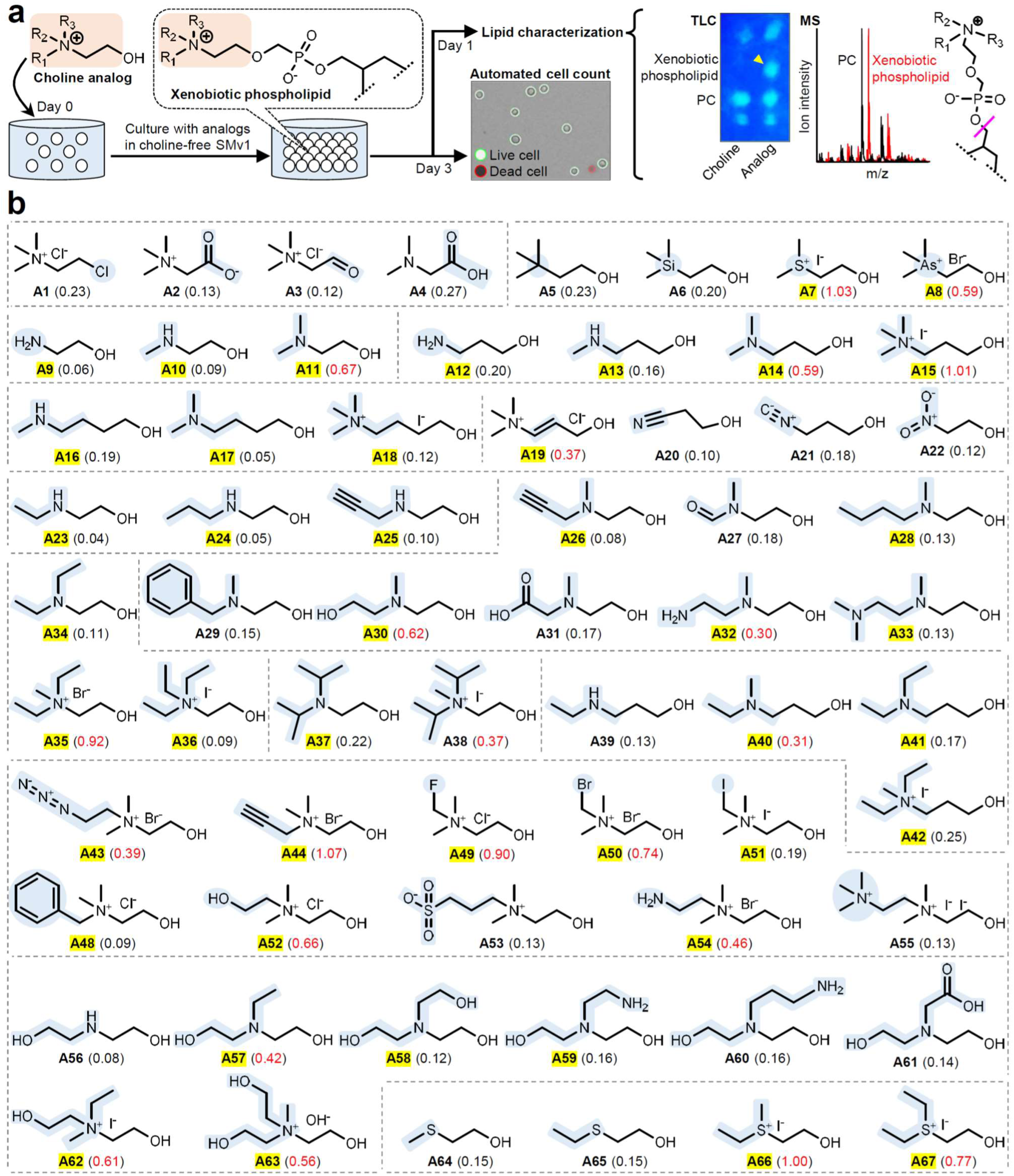
Exploration of choline analogs (part 1). (**a**) Workflow for structure−activity relationship analysis of choline analogs and characterization of lipid products. K562 cells cultured with choline analogs (30 μM) in choline-depleted SMv1 were subjected to the following analyses: (1) automated cell counting to determine proliferation scores relative to the choline-supplemented control, (2) TLC analysis of lipid extract with lipid-sensitive primulin dye to visualize xenobiotic phospholipids, and (3) tandem MS analysis to confirm target xenobiotic phospholipid species by their characteristic ion fragmentation. (**b**) Structures and summarized data of choline analogs. The average proliferation score (n = 3–8 biological replicates) is shown in parentheses, and in red when higher than 0.3. Compound ID is highlighted in yellow when its xenobiotic phospholipids were detected by MS (S/N ratio >10) or TLC. Structural changes of interest in analogs are highlighted in light blue. Tested analogs are grouped by structural features and enclosed by dashed lines. Detailed data are provided in Fig. S1 and Excel file.

Since PC-biosynthetic proteins recognize their substrates through noncovalent interactions such as electrostatic forces^22,23^, amino alcohol derivatives including negative controls were selected. We first examined chloride-substituted analog **A1** and betaine-related metabolites **A2–A4**, all of which lack an alcohol moiety and showed no proliferative activity, confirming that the presence of an alcohol group is essential for activity. Replacing the nitrogen with carbon (**A5**) or silicon (**A6**) abolished activity, while cationic sulfocholine (**A7**) and arsenocholine (**A8**) remained active, underscoring the importance of a positively charged center. TLC and MS analyses showed biosynthesis of **A7**- and **A8**-incorporated phospholipids (Fig. 2b, S1a, S1b). These data demonstrate that xenobiotic phospholipids with unnatural headgroups can support cell proliferation as an alternative to PC.

### Structural differences in phospholipid headgroups impact cellular activities

Biological differences between PC and PE result from the number of *N*-methyl groups in headgroup moieties (Fig. 1a), but their structure–activity relationship remains poorly characterized. We tested **A9–A11** with variable *N*-methyl groups. Although **A9–A11**-incorporated phospholipids were detected, only 2-(dimethylamino)ethanol (**A11**) showed activity. Of **A12–A18** having longer linkers between the nitrogen and the hydroxy group, robust activities were detected in dimethyl and trimethyl aminopropanol derivatives (**A14** and **A15**). Aminopropanol-based analogs containing unsaturated bonds (**A19–A21**), 2-nitroethanol (**A22**), and secondary amine-containing **A23–A25** showed no or lower activity (<0.4). These data suggest that xenobiotic headgroups do not always contribute to proliferation as a substitute for the PC headgroup, and that at least three hydrocarbons surrounding the central cation are required for activity.

We then focused on tertiary amine-containing analogs. Of **A11**-based analogs **A26–A33** where an *N*-methyl group was substituted with various functional groups, the hydroxy- and amino-substituted derivatives remained active (**A30** and **A32**). In contrast to **A11**, its *N*,*N*-diethyl analog **A34** lost activity. Interestingly, **A35** (*N*-methylated **A34**) showed robust activity, but **A36** (*N*-ethylated **A34**) did not. We shifted to quaternary ammonium-containing analogs. Although azidoethylcholine (**A43**) and propargylcholine (**A44**) undergo similar metabolic processing^24,25^, **A44** showed choline-like activity higher than that of **A43**, suggesting that the two headgroups differentially affect cellular functions. Fluoromethylcholine (**A49**) and bromomethylcholine (**A50**) showed robust activities, but iodomethylcholine (**A51**) did not. Of four hydrophilic analogs **A52–A55**, hydroxyethylcholine (**A52**) and aminoethylcholine (**A54**) showed activity. Upon further testing hydroxyethyl-substituted analogs, we also found activities in **A57**, **A62** and **A63**. These observations support the notion that hydrophilic headgroup modifications are tolerable, yet in a structure-dependent manner.

Having observed that only slight differences between *N*-methyl and *N*-ethyl groups frequently produced distinct cellular activities (e.g., **A11** vs **A34**, **A34** vs **A35**), we examined whether this was also the case for sulfur-substituted analogs. No activity was observed in **A64** and **A65**, similar to their corresponding amine analogs **A10** and **A23**. Unexpectedly, *S*-monoethyl- and *S*,*S*-diethyl-substituted sulfocholine (**A66** and **A67**) showed robust activity, in contrast to **A34**, highlighting that even a one-atom difference in headgroup chemistry can impact cellular activities.

### Expanding the chemical diversity of artificial headgroups

For the potential diversity of xenobiotic headgroups, we comprehensively explored chemical modifications in carbon skeletons of amino alcohol derivatives (Fig. 3). We tested **A11**/**A14**-based analogs **B1–B12** which have branched carbon chains between amino and hydroxy groups. Methyl and hydroxymethyl substitutions on the α-carbon were relatively tolerable compared to those on the β-carbon. We extended our analysis to heterocycles, beginning with **A11**/**A14**-based analogs **B13–B21** (three- to five-membered rings). **B16–B19** having a pyrrolidine linker between the methylated amine and the hydroxy group showed activities, but the azetidine counterpart **B14** did not. Interestingly, **B20**, a pyrrolidine analog of non-proliferative **A34**, did not cause lethality. Through extensive analysis of pyrrolidine analogs **B22–B49**, we found that the enantiomers of racemic **B18** (**B34** and **B35**) had different activities from each other. This effect was preserved in the *N*-methyl substituents (**B34M** and **B35M**). Compared with five-membered rings, six-membered ones **B50–B64** showed lower or no activities (score: 0.1–0.4). Unexpectedly, **B65** (*N*-methylated **B55**) and its pyridinium substituent **B69** showed robust activities (score: 0.7–0.8). We additionally designed ester analog **B110** to present carboxylate-containing headgroups via expected enzymatic hydrolysis^26^ (Fig. 3, bottom). MS analysis exhibited both ester and carboxylate forms of **B110**-incorporated phospholipids (Fig. S1b).

**Figure 3.**
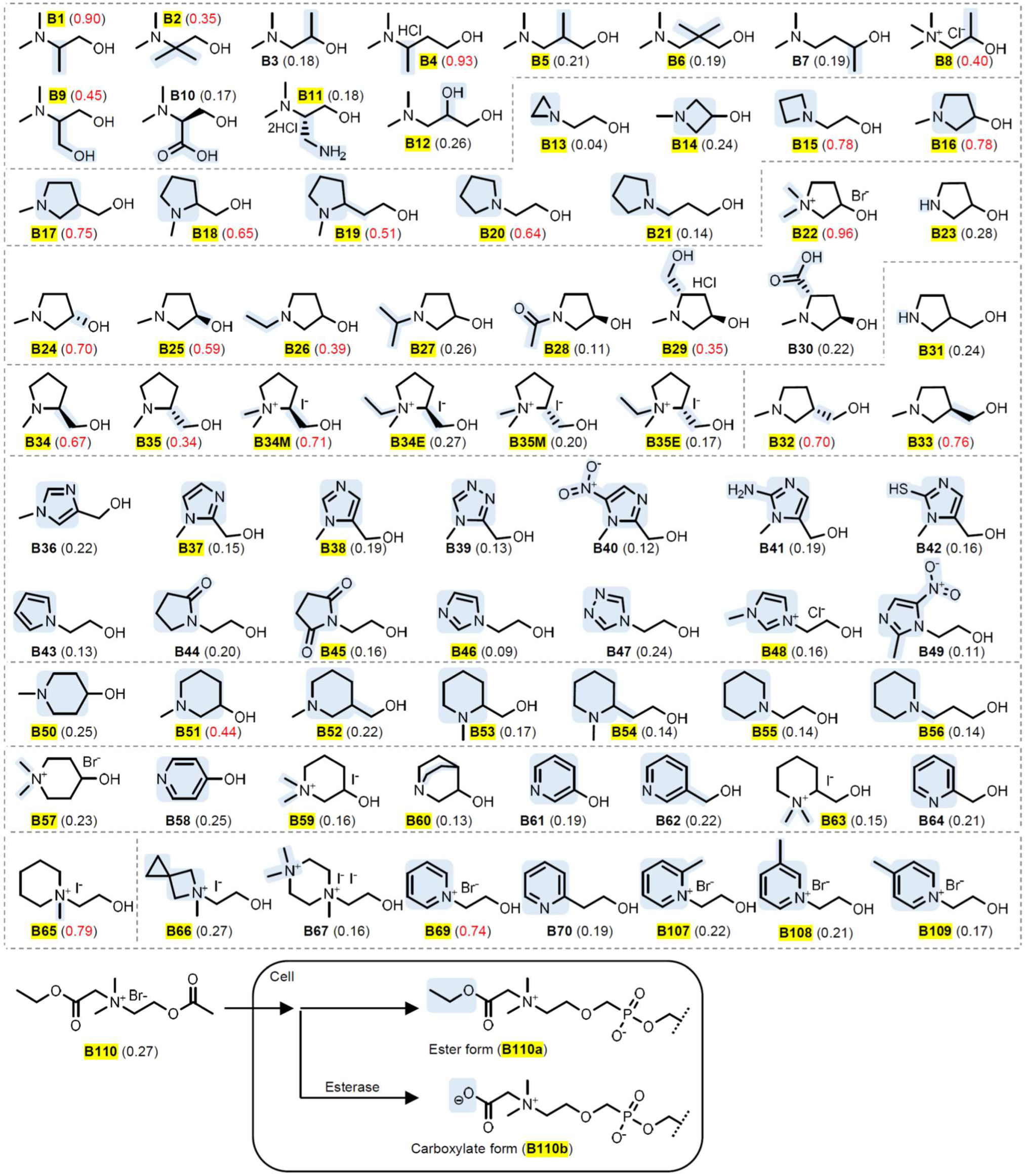
Exploration of choline analogs (part 2). Structures and summarized data of choline analogs. Ester analog **B110** was designed for presenting carboxylate moiety-containing xenobiotic phospholipids through expected enzymatic cleavage inside cells. The average proliferation score (n = 4–6 biological replicates) is shown in parentheses, and in red when higher than 0.3. Compound ID is highlighted in yellow when its xenobiotic phospholipids were detected by MS (S/N ratio >10) or TLC. Structural changes of interest in analogs are highlighted in light blue. Tested analogs are grouped by structural features and enclosed by dashed lines. Detailed data are provided in Fig. S1 and Excel file.

Of the tested 142 compounds, we identified over 100 xenobiotics as headgroup precursors (the precise number depends on the compound concentration and cutoff criteria chosen).

### Effects of xenobiotic phospholipids depend on headgroup structure rather than product yield

To examine the scalability of xenobiotic phospholipid biosynthesis, **A44** was selected because of its proliferative activity and chemical characteristics distinct from choline. TLC analysis showed that **A44**-incorporated phospholipids reached about one-third of total lipids at day 1, and more than half at day 3 with a significant decrease in endogenous PC (Fig. S1a, S1c). This observation demonstrates that our strategy enables predominant production of xenobiotic phospholipids in mammalian cells.

Throughout the TLC and MS analyses, we observed that production levels of xenobiotic phospholipids varied depending on the analog structures, probably due to different affinities to the Kennedy pathway or related transporters. However, there seemed to be no obvious correlation between the production levels and proliferation activities (Fig. S1d). For example, **A23**, **A34** and **A43** showed clear TLC spots and high MS signals compared to **B15**, but no or less activities (Fig. 2b, 3, S1a, S1b), indicating the importance of headgroup structure rather than product yield. Furthermore, distinct activity changes were often linked to slight structural differences, despite similar production levels (e.g., **A11** vs **A34**, **B34** vs **B35**). Additionally, non-proliferative effects were still unchanged under a higher analog concentration (Fig. S1e), but were competitively counteracted by supplementation with choline and dialyzed serum that provides PC via non-Kennedy pathways^17–19,27^ (Fig. S1f), minimizing the possibility of off-target toxicity caused by factors other than xenobiotic phospholipids.

### Xenobiotic phospholipids can serve as membrane lipids

In the Kennedy pathway, the phosphocholine moiety is transferred to diacylglycerol to yield PC, which facilitates bilayer formation^2^ (Fig. 4a, red arrow). When choline is insufficient, neutral lipids instead accumulate to form lipid droplets^2^ (Fig. 4a, green arrow). To determine whether analogs could fulfill the substrate demand, we analyzed cellular metabolic state with LipiORDER dye that distinguishes phospholipid membranes from lipid droplets by ratiometric fluorescence measurements^28^ (Fig. 4a).

**Figure 4.**
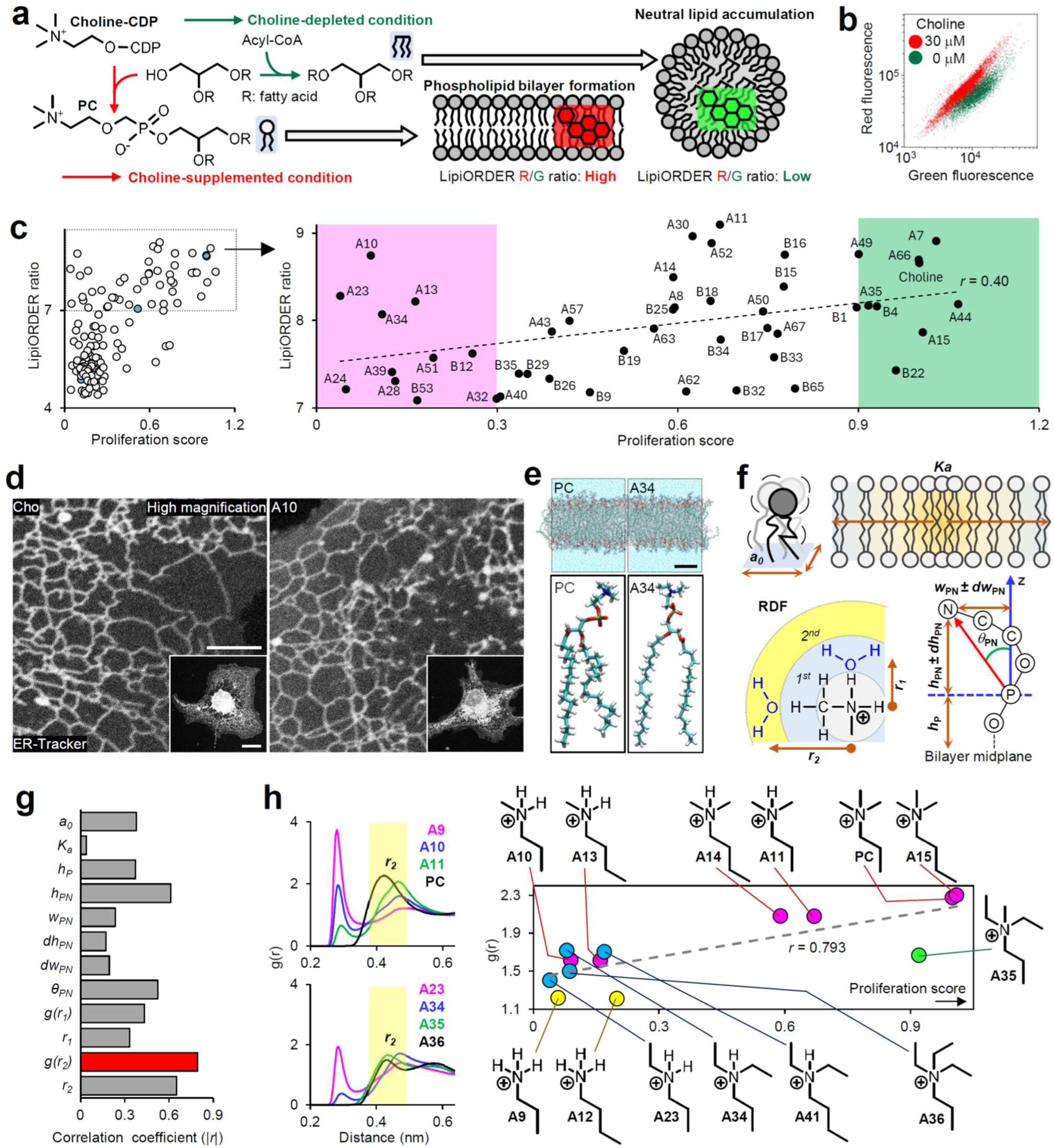
Characterization of xenobiotic phospholipid metabolism. (**a**) Analysis of phospholipid headgroup-forming activity with LipiORDER. (**b**) LipiORDER-based flowcytometric discrimination of metabolic states between choline-supplemented and -depleted K562 cells (detailed data in Fig. S2a and S2b). (**c**) Correlation analysis between LipiORDER ratios and proliferation scores. The dot plot represents the LipiORDER ratio and proliferation score of each analog (detailed data in Fig. S2d). In the left plot, choline controls and analog samples are shown in blue (left to right: 0, 2, 30 μM) and grey, respectively. In the right plot, non-proliferative and proliferative analogs are highlighted in pink and green, respectively. (**d**) ER structures in **A10**-treated COS-7 cells. (**e**) MD simulations for phospholipid bilayer structures (PC and **A34**-modified phospholipids shown as representative). (**f**) The following bilayer properties were calculated: area per lipid *a_0_*; area compressibility modulus *K_a_*; RDF parameters of the peak position *r*_1_*, r*_2_ and peak height *g*(*r*_1_), *g*(*r*_2_); structural parameters *h_P_*, *h_PN_*, *dh_PN_*, *w_PN_*, *dw_PN_*, and *θ_PN_*. (**g**) Correlation analysis between bilayer properties and proliferation scores across 13 phospholipids (detailed data in Fig. S5c). (**h**) Correlation analysis between the RDF parameters and proliferation scores. Left graphs of RDF curves show the distribution probability of water oxygen approaching the nitrogen center of headgroups (the second peak position highlighted in yellow). The dot plot represents the second peak height *g*(*r*_2_) (Fig. S5a) and proliferation score (Fig. 2b) of each phospholipid. The *r* values were calculated by linear regression analysis. Scale bars: **d** (inset), 20 μm; **d** (high magnification), 20 μm; **e**, 2 nm.

Multicolor flowcytometric analysis confirmed that choline depletion decreased the red/green fluorescence ratio – similar to fatty-acid overload – indicating neutral lipid accumulation (Fig. 4b, S2a, S2b). Lipid droplet imaging supported this observation (Fig. S2c). Extending the analysis to analog-treated cells, we compared LipiORDER ratios with proliferation scores (Fig. 4c, S2d). Among the analogs tested, 47 displayed LipiORDER ratios >7.0, exceeding that of the choline-depleted control (ratio: 5.1) (Fig. 4c, enlarged plot; S2b) and suggesting choline-like contributions to headgroup formation. These analogs included both proliferative and non-proliferative types (Fig. 4c, colored areas), showing no correlation between LipiORDER ratios and proliferation scores. When 56 low-ratio analogs (<7.0) were re-examined at a higher concentration, 17 analogs achieved LipiORDER ratios > 7.0 with different proliferation scores (0.1–1.1) (Fig. S2d, S2e, S3). Collectively, these results demonstrate that many analogs can sufficiently consume metabolic intermediates of the Kennedy pathway for headgroup conversions, thereby allowing their structure-dependent differential effects.

Because PC is biosynthesized mainly in the endoplasmic reticulum^1^ (ER), we characterized ER membranes containing xenobiotic phospholipids. Non-proliferative analogs with high LipiORDER ratios (Fig. 4c, pink) (e.g., **A10**) were examined by live-cell ER imaging. ER tracker staining revealed no morphological changes between choline-supplemented and analog-treated K562 cells (Fig. S2f). We then adapted our method to conventional adherent cell lines, establishing culture conditions suitable for assessing the cellular responses to xenobiotic phospholipids (Fig. S4a–S4e). Similar results were obtained in COS-7 cells and HeLa cells, both of which retained normal ER networks under analog treatment (Fig. 4d, S4f, S4g). We further performed direct imaging of alkyne-containing phospholipids through click chemistry^24^. Similar to cells treated with proliferative **A44**, non-proliferative **A26** provided clear fluorescence signals in typical ER networks (Fig. S4h). Given that cellular membranes have lipids with large headgroups including sugar chains and protein conjugation^2^, our results support that xenobiotic phospholipids with sufficient headgroup-forming capacity (i.e. LipiORDER ratio >7.0) can integrate into cellular membranes.

### Xenobiotic headgroups differentially modulate membrane functions

To gain insight into how headgroup structural differences affect membrane properties, we performed molecular dynamics (MD) simulations on 13 representative headgroups bearing variable *N*-methyl and *N*-ethyl substitutions (PC, **A9–A15**, **A23**, **A34–A36**, **A41**). Model phospholipid bilayers were analyzed for 12 typical physical parameters describing lipid geometry, membrane thickness, area compressibility, and radial distribution functions (RDFs) (Fig. 4e, 4f, S5a). Headgroup modifications differentially altered these parameters by −47% to +43% relative to PC (Fig. S5b). Correlation analysis between simulated properties and proliferation scores revealed the strongest relationship for the RDF between the α-carbon of *N*-alkyl groups and water oxygen (NCH···O) (Fig. 4g, 4h, S5c). This correlation was particularly high (*r* = 0.917) among all analogs varying in *N*-methyl groups (PC, **A9–A15**). Interestingly, **A35** deviated from this trend despite having physical parameters similar to those of **A34** (Fig. 4h, S5a, S5b). **A35** differs from **A34** only by the additional *N*-methyl group, which impacts the ability to form a hydrogen bond between the central nitrogen and a proton donor^29^. These findings raise the interesting possibility that artificial headgroup modifications can modulate both physical bilayer properties and phospholipid–protein interactions.

We explored the biological effects elicited by xenobiotic phospholipids. Biosynthesized phospholipids undergo de-acylation and re-acylation remodeling (Fig. 5a), in which various fatty acids are re-incorporated by lysophospholipid acyltransferases with different substrate specificities for headgroup moieties^2^. To determine whether headgroup modifications influence these substrate recognitions, we analyzed the acyl chain compositions by MS profiling of representative phospholipid species (Fig. S6a). Principal component analysis (PCA) uncovered that acyl chain patterns varied depending on headgroup structures (Fig. 5b). Several artificial headgroups displayed acyl chain profiles distinct from deuterium-labeled PC and PE controls (Fig. 5b, pie charts). Altered endogenous phospholipid composition (PE/PC) was also detected (Fig. 5c, S6b). These data prompted us to examine the potential effects of xenobiotic phospholipids on protein activities. The PIEZO1 channel can intrinsically respond to membrane lipid changes^30^. Using fura2 imaging, we found that PIEZO1-mediated Ca^2+^ influx was attenuated in cells cultured with specific analogs (Fig. 5d, 5e, S6c). We further measured Ca^2+^ release triggered by ATP and thapsigargin, which act through different mechanisms^31^. Distinct inhibitory patterns were observed across the tested analogs (Fig. S6d, S6e). PCA showed that the selectivity and potency of inhibition varied with headgroup structures (Fig. 5f), highlighting that xenobiotic phospholipids can differentially influence Ca^2+^ dynamics.

**Figure 5.**
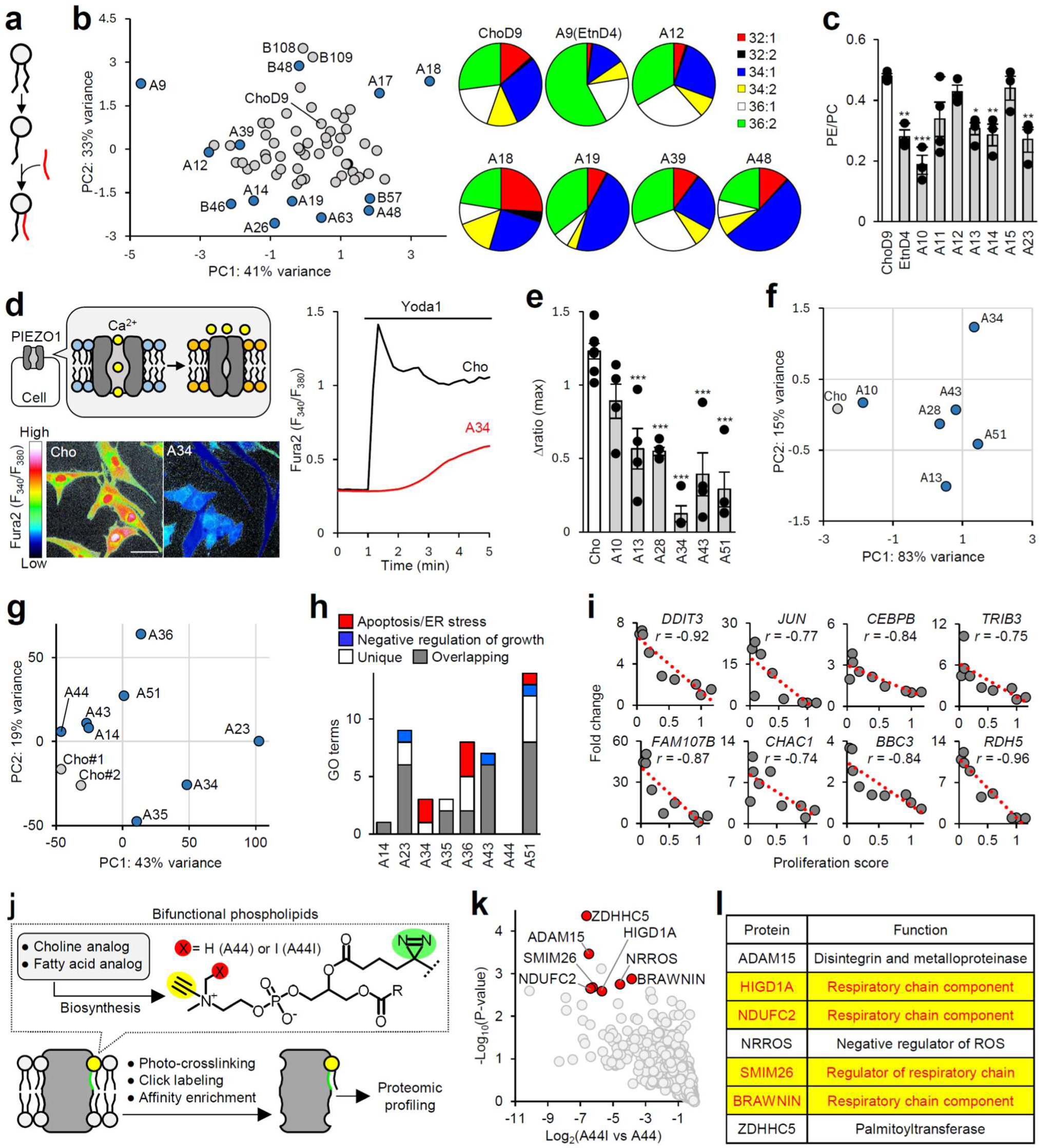
Differential biological effects of xenobiotic headgroups. (**a**) Schematic of phospholipid remodeling. (**b**) PCA of acyl chain compositions. Blue dots represent acyl chain patterns of xenobiotic phospholipids distinct from that of deuterated PC control (choD9). Representative examples are shown as pie charts. (**c**) Altered endogenous lipid composition in analog-treated K562 cells (mean ± SEM, n = 3–4 biological replicates). (**d, e**) Inhibition of PIEZO1-mediated Ca^2+^ influx in analog-treated C2C12 cells. Representative images and traces are shown (**d**). The Ca^2+^ influx was quantified as the maximal increment of fura2 ratio (mean ± SEM, n = 4–7 biological replicates) (**e**). (**f**) PCA of Yoda1-, ATP- and thapsigargin-induced Ca^2+^ responses (detailed data in Fig. S6c–S6e). (**g**) PCA of transcriptome responses in analog-treated K562 cells. (**h**) GO enrichment analysis (analogs vs choline). The GO terms of biological processes were identified as statistically significant (|log_2_(fold change)| >1, FDR <0.05, adjusted *p* value <0.05). (**i**) Correlation analysis between upregulated transcripts (normalized FPKM values) and proliferation scores. The *r* values were calculated by linear regression analysis. (**j**) Workflow for phospholipid−protein interactome analysis. (**k**) Volcano plot depicts comparison of phospholipid-linked proteins in (**j**). Red dots represent the top-ranked integral membrane proteins that satisfy the following criteria: log_2_(fold change of **A44I** vs **A44**) <−3.0, −log_10_(*p*-value) >2.5, FDR <0.01. (**l**) Table shows significantly enriched proteins in (**k**). Mitochondrial proteins are highlighted. \**p* <0.05, \*\**p* <0.01, \*\*\**p* <0.001 (one-way ANOVA). Scale bar: **d**, 50 μm. Detailed data (**b**, **h**) are provided in Excel file.

### Identification of headgroup structure-sensitive membrane proteins

To investigate broader cell responses to headgroup alterations, eight analogs with varying proliferation scores were subjected to RNA-seq analysis under the 500 μM condition where all of the analogs achieved LipiORDER ratios >7.0 (Fig. S3). PCA and gene ontology (GO) enrichment of differentially expressed genes relative to the choline-supplemented control revealed distinct transcriptomic signatures among analog-treated cells (Fig. 5g, 5h). Notably, analogs with proliferation scores <0.4 provided GO terms related to “negative regulation of growth” and “intrinsic apoptotic signaling pathway in response to ER stress” (Fig. 5h). Consistent with this, several genes associated with stress responses^32^ and regulated cell death^33^ exhibited inverse correlations between expression levels and proliferation scores (Fig. 5i). As found in the cases of PC vs **A51** and **A34** vs **A35**, headgroup modifications on the order of one atom made a significant difference in signaling cascades.

We sought to define the basis for the observed vulnerability to such headgroup modifications. We set up headgroup-focused phospholipid–protein interactome analysis with biosynthesized bifunctional phospholipids which have both clickable alkyne and photoactivatable diazirine^27^ (Fig. 5j). Given the activities of PC-like **A44** and non-PC-like **A51** (Fig. 2b, 5h), we prepared iodomethyl-substituted propargylcholine (**A44I**) (Fig. S6f), expecting altered protein−headgroup recognition^34^. We confirmed the biological and chemical characteristics of **A44I**-incorporated phospholipids (Fig. S6g–S6i). Through quantitative comparison of phospholipid-linked proteomes (Fig. 5j), the detected 3,926 proteins were narrowed down to the top-ranked seven integral membrane proteins (0.2%) that were significantly more enriched in **A44**-treated samples than in **A44I**-treated ones (Fig. 5k, red dots). Notably, four of these are functionally associated with mitochondrial respiratory chain complexes^35–39^ (Fig. 5l). In line with this, iodinated PC (**A51**) led to mitochondrial depolarization (Fig. S6j), which agrees with previous studies of mitochondrial abnormalities caused by PC metabolic defects^2,7,10,18,40,41^. These data demonstrate that our structure–activity relationship-based strategy enables identification of phospholipid–protein interactions sensitive to subtle headgroup structural alterations.

### Extension of xenobiotic phospholipid rewiring to primary cells

To explore the applicability of this method, we focused on primary cells with specialized functions. Differentiated heart muscle cells, cardiomyocytes, possess deeply invaginated membranes known as T-tubules, which are critical for Ca^2+^-dependent excitation–contraction coupling^42^. Given the active PC biosynthesis^43,44^, we examined T-tubule membrane structures under different xenobiotic conditions. Adult rat ventricular cardiomyocytes were cultured with selected analogs in choline-free M199 medium, followed by staining T-tubules with FM4-64 dye (Fig. S4a). Choline- and **A15**-treated cardiomyocytes retained the characteristic striped pattern of T-tubules, comparable to freshly isolated controls, without toxicity or abnormal beating (Fig. 6a, S4i, S4j). In contrast, **A10** and **A13** caused severe morphological abnormalities with the majority of cells adopting a rounded shape (Fig. S4i), which agrees with the non-proliferative effects on cell lines (Fig. 2b, S4b-S4d). Among the few surviving cells that preserved their elongated morphology, abnormal spontaneous contractile activity was observed. Notably, **A10** induced extensive membrane blebbing, small aggregates, wave-like propagation, fibrillation-like beating and hypercontraction events despite no external electrical stimulation (Fig. 6a, Movie S1). **A13**-treated cells occasionally exhibited irregular spontaneous contractions without prominent membrane abnormalities (Fig. S4j, Movie S2).

**Figure 6.**
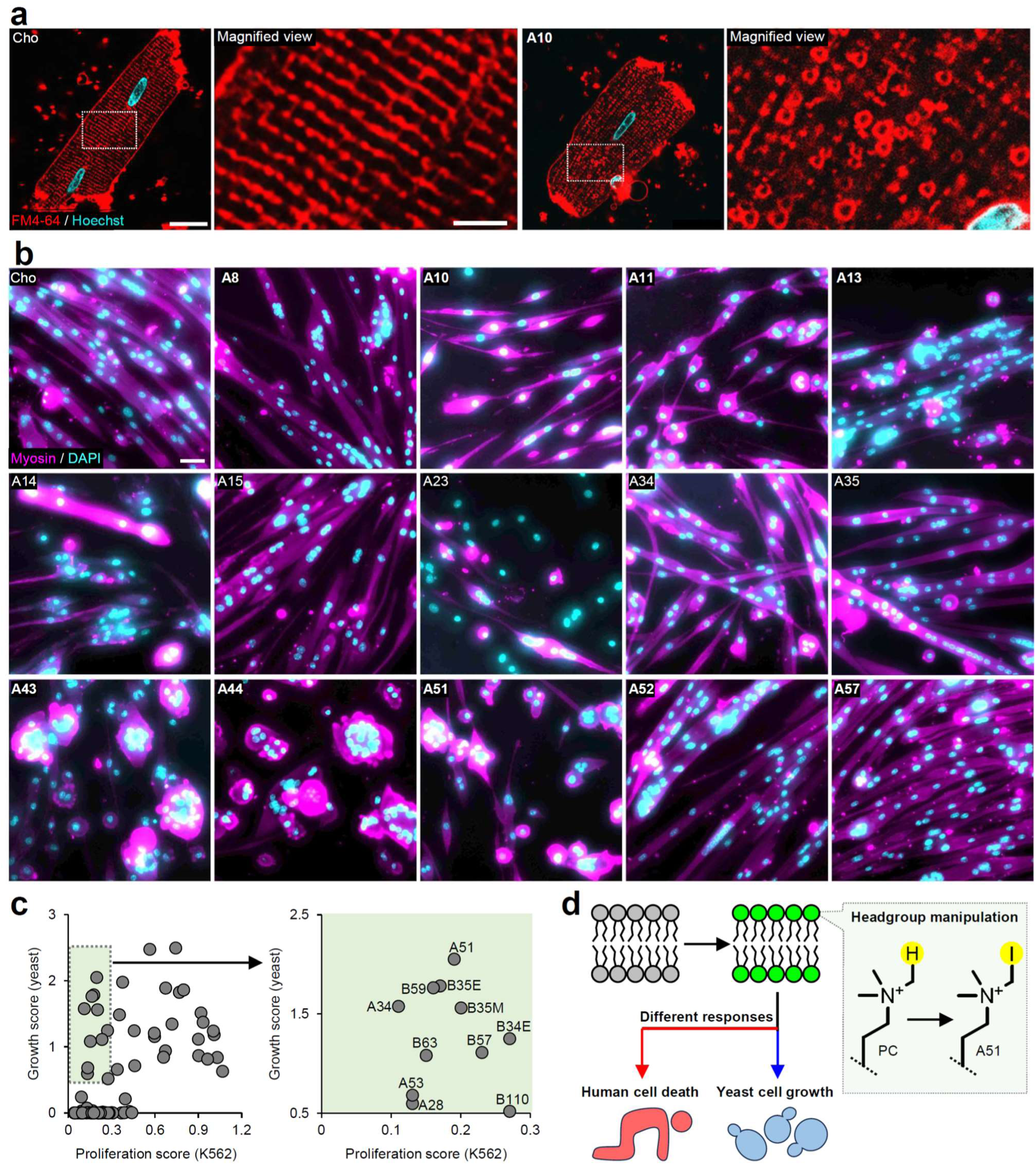
Applications in cardiomyocytes, skeletal muscle stem cells and yeast. (**a**) Disorganized T-tubules in **A10**-treated cardiomyocytes. Cardiomyocytes isolated from rat ventricles were cultured with choline and **A10** (1 mM) in M199 media for 4 hours, and stained with FM4-64 and Hoechst 33342 for live-cell confocal microscopy. (**b**) Diverse morphological phenotypes in analog-treated muscle stem cells. Skeletal muscle stem cells isolated from mouse limb (Fig. S4k) were differentiated with choline and analogs (500 μM) in B27-supplemented DMEM for four days, and immunostained against muscle-specific myosin heavy chain with nuclear staining for epifluorescence microscopy. (**c**) Cross-species comparison of cellular activities of xenobiotic phospholipids. Yeast *cho2Δopi3Δ* cells were cultured with analogs (100 μM) in choline-free SD medium for two days, and subjected to ATP measurements to determine their growth scores compared to the choline-supplemented control (n = 2–3 biological replicates). The dot plot represents the average yeast growth score and the K562 cell proliferation score (Fig. 2b and 3) of each analog. Green color highlights analogs that have non-proliferative effects on K562 cells (score <0.3) but growth effects on yeast cells (score >0.5). (**d**) Model representing different phospholipid headgroup preferences between human and yeast cells. As a case example, iodination of the PC headgroup impacts human cells but not yeast. Scale bars: **a**, 20 µm; **a** (magnified view), 5 µm; **b**, 20 µm.

We turned to a differentiation model of skeletal muscle stem cells, in which myogenic cells fuse to form myotubes through dynamic membrane reorganization^45,46^. Mouse muscle stem cells were differentiated with selected analogs in choline-free DMEM (Fig. S4a, S4k). Immunostaining of muscle-specific myosin revealed diverse morphological phenotypes depending on xenobiotic headgroup structures (Fig. 6b): typical elongated myotubes, impaired cell fusion, and rounded syncytia. Interestingly, we found examples of analogs whose effects were uncorrelated between muscle cells and K562 cells: myotube formation was disturbed by proliferative **A44**, but facilitated by less proliferative **A57** (Fig. 2b, 6b). Together, these results demonstrate that xenobiotic phospholipids can modulate specialized cellular functions and differentiation programs.

### Different headgroup preferences between human and yeast cells

Lastly, we extended this method to a non-mammalian model organism. Given that yeast *S. cerevisiae* can grow without PC under specific conditions^9^, we were interested in whether essential phospholipid headgroups differ between human and yeast. Because the yeast *cho2Δopi3Δ* mutant requires exogenous choline for Kennedy pathway-mediated PC biosynthesis and cell growth^9^, we cultured *cho2Δopi3Δ* cells in synthetic defined (SD) medium containing choline or selected analogs to assess growth responses (Fig. S4l). Remarkably, several non-proliferative analogs in K562 cells promoted the yeast growth at levels comparable to that of the choline-supplemented control (Fig. 6c, green area), indicating that yeast and human cells differ in their response to headgroup modifications. As Ire1 sensor is known to respond to general lipid bilayer stress including defective PC biosynthesis^47^, Ire1 activity was indeed increased in choline-depleted *cho2Δopi3Δ* cells, but minimally detected in choline-and **A51**-treated cells (Fig. S6k). Taken together, these data highlight interspecies differences in headgroup preferences, emphasizing their significance for cell life and death (Fig. 6d).

## Discussion

Our understanding of membrane lipids remains limited due to a lack of robust tools for investigating metabolite structure–activity relationships. Previous studies have successfully introduced clickable handles into the PC headgroup for several applications^15,16,24,25,27,43^. However, the potential for large-scale production of artificial headgroups as intervention tools for cellular functions has been poorly explored. Here, we developed a metabolic rewiring method to manipulate phospholipid headgroups and demonstrated efficient biosynthesis of xenobiotic phospholipids (Fig. 1–3). These products accounted for a substantial proportion of total lipids, exceeding levels previously reported^13,14^. This outstanding scalability seems to benefit from the energetically favorable reactions of the Kennedy pathway^7^. For expanding headgroup chemistry, we identified over 100 xenobiotics covering various functional groups. This unprecedented diversity broadens the scope of membrane lipid manipulation. Importantly, our strategy circumvents both genetic engineering and total phospholipid synthesis, relying instead on simpler alcohol derivatives as programmable metabolic precursors.

Using this platform, we demonstrated that xenobiotic headgroups elicit diverse and structure-dependent biological effects, providing evidence that headgroup modifications can affect lipid–protein interactions and differentially modulate various cellular processes across cell types and kingdoms (Fig. 4–6). Furthermore, we illustrated that this method can be used as an efficient platform to reveal previously unrecognized roles of phospholipid headgroups. A striking finding is that minor structural changes – such as single atom substitution (e.g., PC vs **A51**) – can shift human cells from proliferation to death. Our data suggest that specific headgroups perceived as non-PC can trigger stress signaling, highlighting a lipid structure-sensitive unique mechanism in mammalian cells. Indeed, we unveiled headgroup-sensitive human membrane proteins (Fig. 5l). The identified four mitochondrial proteins are microproteins (8-14 kDa) that seemingly have intrinsically disordered regions but contribute to respiratory complex assembly and regulation^35–39^. None of these protein sequences is well conserved in yeast, which is in line with our finding that iodinated PC impacts human cells but not yeast (Fig. 6d). Our discovery agrees with previous reports that disease-associated PC abnormalities cause mitochondrial dysfunction^2,7,10,18,40,41^. Overall, these results exemplify the evolutionary diversity of key phospholipid–protein interactions, illustrating why the PC headgroup is significant for humans.

We expect that this method would be applicable to various eukaryotic models, particularly for synthetic medium-based culture systems^20^ (e.g., iPS cells, cell therapies, culture meat). Furthermore, it could potentially be extended to plants and animals under chemically defined conditions^16,24^. Given that nutrient depletion and inhibition of lipid biosynthetic enzymes alter metabolite pools^2^, our method provides a complementary means of dissecting the specific roles of phospholipid headgroups. Besides phospholipid–protein interactome mapping, this method is compatible with genetic screens for comprehensive identification of the key molecules^16,48^. Although this paper focused on the Kennedy pathway, choline analogs might be transformed into other choline-derived metabolites including lysoPC, sphingomyelin and acetylcholine^7^. Beyond modulating metabolite material properties, this approach may open up opportunities for intracellular organic chemistry^49,50^. Finally, we anticipate that this study will facilitate the engineering of eukaryotic membranes to create novel biological systems.

## Acknowledgment

This work was supported by JSPS KAKENHI Grant Numbers JP23H03856, JP25K03034, the University of Shizuoka Academic Research Grant, Kanamori Foundation, Astellas Foundation for Research on Metabolic Disorders, Chugai Foundation for Innovative Drug Discovery Science, and Ono Medical Research Foundation to M.T.; and Intramural Research Grant (5-6) for Neurological and Psychiatric Disorder of NCNP and Takeda Science Foundation (Specific research grant) to Y.H.

## Author contributions

Conceptualization: M.T. Methodology: R.N., K.H., Ak.M., T.K., Y.Uj., Ta.S., To.S., S.M., M.I., Y.H., and M.T. Investigation: R.N., Ai.M., Y.Uc., Y.O., N.A., Y.K., K.H., S.N., Y.Uj., K.S., Ta.S., To.S., M.I., and M.T. Writing – original draft: R.N. and M.T. Writing – review & editing: R.N., K.H., N.T., T.K., Y.Uj., Ta.S., To.S., S.M., M.I., Y.H. and M.T.

## Competing interests

The authors declare no competing interests.

## Methods

### Materials

Reagents and resources used in this study are summarized in Excel file.

### Choline analogs

Commercially available analogs were used without further purification. For in-house preparation of analogs, the synthetic methodology was adapted from the previous report^51^. Method A: To a solution of the amino alcohol derivative (2.0 mmol) in THF (1 mL), the desired alkyl halide (3 equiv.) was added. The reaction mixture was stirred at 60°C for 6 h. After completion, the THF solution was decanted from the crude solid, which was subsequently washed with Et_2_O (3 × 10 mL) to afford the corresponding quaternary ammonium halide as a colorless solid. Method B: To a solution of the amino alcohol derivative (2.0 mmol) in THF (1 mL), the desired alkyl halide (3 equiv.) was added. The reaction mixture was stirred at 60°C for 6 h. The THF solution was decanted from the residue, which was then dissolved in MeOH (1.0 mL). Et_2_O was added until phase separation occurred. After brief sonication, the upper layer was removed. This dissolution and washing procedures were repeated three times to afford the quaternary ammonium halide as a pale-yellow viscous compound. For lipid analysis by MS, deuterium-labeled choline (choD9) and ethanolamine (etnD4) were used. Detailed information and characterization data of analogs are summarized in Excel file.

### Cell lines

For usual cell maintenance with conventional serum-containing media, K562 cells were cultured in IMDM supplemented with 1% Penicillin–Streptomycin and 10% FBS at 37°C with 5% CO_2_. HeLa, COS-7, and C2C12 cells were cultured in DMEM supplemented with 1× Penicillin–Streptomycin and 10% FBS at 37°C with 5% CO_2_. For serum-free experiments, cells were cultured in synthetic media as shown in Fig. S4a. Detailed information is provided in Excel file.

K562 cells were subjected to proliferation assay, LipiORDER analysis, ER imaging, lipid droplet imaging, lipid analysis, mitochondria membrane potential assay, transcriptomics and proteomics. HeLa cells were subjected to ER imaging and Ca^2+^ imaging. COS-7 cells were subjected to ER imaging and phospholipid imaging. C2C12 cells were subjected to Ca^2+^ imaging.

### Proliferation assay

K562 cells were seeded into 12-well plates (1 mL of 5 × 10^4^ cells/mL) and cultured with 30 or 500 μM choline analogs for three days in choline-depleted SMv1. Alternatively, K562 cells were cultured with 30 µM choline metabolites or 10 µM lysophospholipids in choline-depleted SMv1. For inhibition of the Kennedy pathway, serial dilutions (0.001 to 3 μM) of RSM932A were added to SMv1. For evaluation of proliferation under vitamin deficiency conditions, K562 cells were cultured for three days in SMv1 individually lacking one of the ten vitamins. For rescue experiments, choline (500 μM) or dialyzed FBS (5%) was added to analog-containing SMv1. Cell cultures were treated with trypan blue for measurements of cell numbers and live cell percentages with an automated cell counter (Countess, Thermo Fisher Scientific). Proliferation scores of cells treated with analogs were determined by live cell numbers relative to 30 μM choline-supplemented control.

### LipiORDER analysis

K562 cells were seeded into 12-well plates and cultured for 1 day in SMv1 containing 30 or 500 μM choline or analogs. The cell cultures were incubated with 500 nM LipiORDER at 37°C for 10 min and analyzed by flow cytometry (MA900, Sony). Instrument settings were used as previously described^52^. For each sample, at least 10,000 events were analyzed. The data acquisition, analysis, and image preparation were carried out using the instrument software MA900 Cell Sorter Software (Sony). Representative gating strategies were presented in Fig. S2a.

### Mitochondria membrane potential assay

K562 cells were seeded into 12-well plates and cultured for 3 days in SMv1 containing 500 μM choline or **A51**. The cell cultures were incubated with 10 nM tetramethylrhodamine ethyl ester (TMRE) at 37°C for 30 min and analyzed by flow cytometry (MA900, Sony). For each sample, at least 30,000 events were analyzed. The data acquisition, analysis, and image preparation were carried out using the instrument software MA900 Cell Sorter Software (Sony).

### Confocal imaging

For ER labeling, K562, COS-7, and HeLa cells were cultured with 500 μM choline or analogs in synthetic media (Fig. S4a) for 2−4 days, and stained with ER-Tracker Red (1:1000) at 37°C for 10 min. For lipid droplet labeling, K562 cells were cultured with 500 μM choline or analogs in SMv1 (Fig. S4a) for 1 day, and stained with LipiDye^®^Ⅱ (1:1000) at 37°C for 10 min. For phospholipid labeling, COS-7 cells were cultured with 30 μM analogs in the synthetic medium (Fig. S4a) for 1 day, fixed with 4% paraformaldehyde at room temperature for 60 min. After washing with 1% BSA/PBS, alkyne-containing phospholipids were labeled with Alexa Fluor 488 azide using Click-iT kit. Cells labeled with fluorescent dyes were analyzed by a Zeiss LSM800 confocal microscope equipped with a 63× oil-immersion objective. The image preparation was carried out using the instrument software ZEN (ZEISS).

### Lipid analysis

K562 cells were cultured with 30 μM choline or analogs in SMv1 for 1 day, and analyzed by an automated cell counter to collect the same lipid amount among tested samples. Cell pellets corresponding to 20 μg of total protein were subjected to total lipid extraction by the Bligh & Dyer method. For TLC analysis, extracted lipids were separated on a TLC plate (Silicagel 70 TLC Plate-Wako) with the solvent system of chloroform/methanol/water (65:25:4, v/v/v), stained by spraying with a primuline solution (0.001% in 80% acetone), and visualized under UV illumination. For MS analysis, extracted lipids were analyzed as previously described^53^, using an LC-MS system consisting of a triple quadrupole mass spectrometer (LCMS-8040, Shimadzu) coupled to a pump (LC-30AD, Shimadzu), an autosampler (SIL-30AC, Shimadzu), and a column oven (CTO-20AC, Shimadzu). LC-MS/MS analysis was performed using an Accucore RP-MS column (2.6 μm, 2.1 mm × 50 mm; Thermo

Fisher Scientific, Waltham, MA) equipped with a SecurityGuard C18 guard column (Phenomenex, Torrance, CA). Mobile phase A consisted of H_2_O/acetonitrile (60:40, v/v), and mobile phase B consisted of isopropanol/acetonitrile (90:10, v/v). Mobile phases A and B were supplemented with 10 mM ammonium formate and 0.1% formic acid. The gradient elution program was set as follows: B% = 10–10–100–10–10 (0–2–15–15.01–20 min). The column temperature was maintained at 40°C, and the flow rate was 0.4 mL/min. Mass spectrometry was conducted with the following parameters: nebulizer gas flow, 3.0 L/min; drying gas flow, 15.0 L/min; desolvation line temperature, 250°C; heat block temperature, 400°C and collision-induced dissociation gas pressure, 230 kPa. Ion fragmentation patterns of individual molecular species derived from xenobiotic phospholipids and natural phospholipids were analyzed by multiple reaction monitoring. Detailed analytical parameters for xenobiotic phospholipids are provided in Excel file. For comparison of cellular phospholipid composition, authentic standards of each phospholipid (PC, PE, PI, PG, PS) were added to lipid extracts and used as references for data normalization.

### Ca^2+^ imaging

HeLa and C2C12 cells were cultured with 500 μM choline or analogs in synthetic media (Fig. S4a) for 2 and 4 days. Intracellular Ca^2+^ measurements were performed according to the previous reports^54,55^. Cells were loaded with 3 μM Fura2 AM at 37°C for 40 min for monitoring following Ca^2+^ responses. For measuring PIEZO1-mediated Ca^2+^ influx, the extracellular solution was replaced with HEPES-buffered saline (2Ca-HBS; 140 mM NaCl, 5 mM KCl, 2 mM CaCl₂, 2 mM MgCl₂, 10 mM D-glucose, and 10 mM HEPES, pH adjusted to 7.4 with NaOH). After recording the baseline response for 1 min, Yoda1-containing 2Ca-HBS was added to reach a final Yoda1 concentration of 10 μM. The fluorescence was recorded for an additional 4 min. For evaluation of Ca²⁺ release from the ER, cells were immersed with Ca²⁺-free HEPES-buffered saline (0Ca-HBS; 140 mM NaCl, 5 mM KCl, 2 mM MgCl₂, 10 mM D-glucose, 0.5 mM EDTA and 10 mM HEPES, pH adjusted to 7.4 with NaOH). After recording the baseline response for 1 min, 0Ca-HBS containing ATP or thapsigargin was added to reach a final concentration of 10 μM ATP or 2 μM thapsigargin. The response was recorded for an additional 4 min. Time-lapse images were recorded every 10 s. Ratiometric images (F_340_/F_380_) were analyzed with Physiology software (Zeiss). The ratiometric response to Yoda1, ATP and thapsigargin was quantified as the difference between the highest value and the mean during the first 1 min.

### Transcriptome analysis

K562 cells were cultured in SMv1 containing 500 μM choline or analogs for 2 days. Total RNA was extracted using the RNeasy^®^ Micro Kit according to the manufacturer’s instructions. RNA sequencing libraries were prepared using NEBNext Poly(A) mRNA Magnetic Isolation Module and NEBNext Ultra II Directional RNA Library Prep Kit. Samples were sequenced with Illumina NovaSeq X Plus (150bp×2 paired-end, 40.0 M reads per sample). Raw paired-end RNA-seq reads were subjected to quality assessment using FastQC (v0.11.7). Adapter sequences and low-quality bases (Phred score <20) were removed with Trimmomatic (v0.38) using the following parameters: ILLUMINACLIP:path/to/adapter.fa:2:30:10 LEADING:20 TRAILING:20 SLIDINGWINDOW:4:15 MINLEN:36. The trimmed reads were aligned to the reference genome using HISAT2 (v2.1.0). The resulting SAM files were converted to BAM format with Samtools (v1.9). Gene-level read counts were quantified using featureCounts (v1.6.3), and expression levels were normalized as transcripts per million (TPM). For exploratory data analysis, normalized count data were used to perform hierarchical clustering based on Euclidean distances using the stats (v3.6.1) and gplots (v3.0.1.1) R packages. Principal component analysis (PCA) was conducted to visualize sample variance across the first two principal components. After normalization by the trimmed mean of M-values (TMM), differential expression analysis was performed with edgeR (v3.26.8). Genes with an absolute log₂ fold change >1 and a false discovery rate (FDR) <0.05 (Benjamini–Hochberg correction) were considered significantly differentially expressed. Gene Ontology (GO) enrichment analysis of differentially expressed genes (DEGs) was carried out using GOATOOLS (v1.1.6). P values were adjusted using the Benjamini–Hochberg method for multiple calibrations.

### Interactome analysis

K562 cells (2 mL of 1 × 10^6^ cells/mL in a 6-well plate) were cultured with 500 μM analogs together with 100 μM 16-diazFA in SMv1 for 1 day. Cells were collected in 2 mL tubes, washed once with ice-cold PBS, resuspended in 1 mL of ice-cold PBS and transferred to 35 mm dishes. Each sample was photo-crosslinked on ice by UV irradiation at 365 nm for 20 min using a metal cooling block. Cells were collected into 1.5 mL tubes, and the medium was removed. Pellets were resuspended in 300 μL PBS, followed by addition of methanol/chloroform (4:1, v/v; 750 μL). After vortexing, the samples were centrifuged at 13,200 × g for 5 min at 4°C. After the supernatant was removed, the pellets were resuspended in 180 μL of lysis buffer (150 mM NaCl, 50 mM HEPES, 1% TritonX-100 and 1× cOmplete, EDTA-free, pH adjusted to 7.4 with NaOH), and incubated at room temperature for 30 min with gentle tapping every 10 min. For click labeling, 20 μL of 10× reaction mixture (10 mM CuSO4, 20 mM Tris(3-hydroxypropyltriazolylmethyl)amine (THPTA), 10 mM Biotin-PEG3-Azide and 100 mM Sodium L-Ascorbate) was added to the lysate and incubated at 37°C for 30 min with gentle agitation. Proteins were precipitated by adding 800 μL of ice-cold acetone, followed by centrifugation at 5,000 × g for 20 min at 4°C. The pellets were washed with 800 μL of ice-cold methanol and centrifuged again under the same conditions. The resulting protein pellets were air-dried, and analyzed by following SDS-PAGE or LC-MS/MS.

For SDS-PAGE, the protein pellets were resuspended in 100 µL of SDS-PAGE sample buffer. Control samples without UV irradiation were also prepared for comparison. After electrophoresis using a precast gel, proteins were transferred onto a PVDF membrane and blocked with 1.5% BSA in TBS-T for 1 h at room temperature. The membrane was then incubated with horseradish peroxidase (HRP)-conjugated streptavidin (1:4000) for 1 h at room temperature. Labeled proteins were visualized using the West Pico PLUS Chemiluminescent Substrate. Total protein was detected by Coomassie Brilliant Blue (CBB) staining of a parallel gel loaded with the same samples.

For LC-MS/MS, sample processing was performed according to the previous report^56^ with minor modifications. Briefly, the protein pellets were resuspended in 200 µL of 2% SDS-TBS and sonicated using a closed-type ultrasonic disruptor. Protein concentrations were determined using a ProteoAnalyzer (Cat# 5191-6640, Agilent) and adjusted to 1 μg/µL with 2% SDS-TBS. The samples were centrifuged at 15,000 × g for 30 min at room temperature. The pellets were resuspended in 40 µL of Sera-Mag SpeedBead Streptavidin-Blocked Magnetic Particles (Cytiva) and incubated with gentle agitation at 2000 rpm for 1 h. The magnetic beads were washed sequentially with the following buffers: 0.5% SDS–TBS containing 25 mM TCEP, RIPA buffer containing 25 mM TCEP, 2 M urea–TBS-T containing 25 mM TCEP, 1% BSA–TBS-T, 0.5% SDS–TBS containing 25 mM TCEP, RIPA buffer containing 25 mM TCEP, 2 M urea–TBS-T containing 25 mM TCEP, 1 M KCl–0.01% LMNG, 0.1 M Na₂CO₃–0.01% LMNG, and finally 50 mM Tris-HCl (pH 8.0) containing 0.024% LMNG and 10 mM CaCl₂. Each washing step was performed by centrifugation at 3000 rpm (33000 rpm for 1%BSA–TBS-T) for 10 min (1 min for the final wash), followed by removal of the supernatant. The magnetic beads were resuspended in 100 µL of 50 mM Tris-HCl (pH 8.0) containing 0.024% LMNG and 10 mM CaCl₂. Bound proteins were digested with 500 ng Trypsin/Lys-C (Promega) at 37°C for 18 h. The supernatant was collected, reduced and alkylated with 10 mM TCEP and 40 mM 2-chloroacetamide at 80°C for 15 min, acidified with 5% TFA, and desalted using GL-Tip SDB (GL Sciences). Peptides were eluted with 34% acetonitrile containing 0.1% TFA, dried under vacuum, and reconstituted in 10 µL of 0.1% TFA with 0.02% DMNG. Samples were analyzed by nanoLC-MS system. Mobile phase A: 0.1% formic acid and B: 0.1% formic acid in 80% acetonitrile. Mobile phase gradient: 1% B at 0.5 min, 10% B at 4 min, 40% B at 49 min and 98% B at 52 min. The flow rate was 800 nL/min during the initial 0.5 min and the final 1.5 min, and 200 nL/min for the remainder of the run. Peptide eluents were analyzed by Orbitrap Exploris 480 MS (Thermo Fisher Scientific). The acquired data were analyzed using the DIA-based proteomics software DIA-NN to identify and quantify proteins with both precursor and protein FDRs controlled below 1%. Detailed results are provided in Excel file.

### Cardiomyocytes

Cardiomyocyte experiments were conducted in accordance with the *Guide for Animal Experimentation, Nagoya Institute of Technology*. Ventricular cardiomyocytes were isolated from adult male Slc:SD rats using a modified Langendorff perfusion method, as previously described^57^. Briefly, rats were anesthetized with sevoflurane, and their hearts were excised and mounted on a perfusion apparatus via the aorta using an 18 G hypodermic needle (NN-1838S, Terumo). The hearts were perfused retrogradely with a calcium-free cell isolation buffer (CIB; 130 mM NaCl, 5.4 mM KCl, 0.5 mM MgCl₂, 0.33 mM NaH₂PO₄, 22 mM glucose, and 25 mM HEPES, pH 7.4) containing 0.4 mM EDTA to remove blood and chelate extracellular calcium. The perfusate was then replaced with an enzyme solution (ES; CIB containing 1 mg/mL collagenase, 0.06 mg/mL trypsin, and 0.06 mg/mL protease) supplemented with 0.3 mM CaCl₂. After digestion, the ventricles were dissected and minced into small pieces, which were gently agitated in the enzyme solution containing 0.7 mM CaCl₂ and 2 mg/mL BSA at 37°C. The dissociated cells were collected by centrifugation and gently resuspended in CIB containing 1.2 mM CaCl₂ and 2 mg/mL BSA. The suspension was incubated at 37°C for 10 min. After incubation, the cells were centrifuged again, and most of the supernatant was discarded. The remaining small volume of the solution was used to gently resuspend the cell pellet, and the resulting suspension was transferred onto glass-bottom dishes (D11530H, Matsunami) that had been pre-coated with laminin and prepared with choline-free Medium 199. The cells were allowed to adhere to the bottom surface by incubation at 37°C for 30 min. After attachment, the medium was replaced with choline-free Medium 199 supplemented with 1 mM choline or analogs. The cells were then incubated for 4 h at 37°C with 5% CO₂ and observed using an inverted microscope (IX73, Olympus).

For fluorescence observation, cardiomyocytes cultured under each condition were gently washed and incubated with 1.4 μg/mL FM4-64 and 4 μg/mL Hoechst 33342 diluted in the same choline-free Medium 199 formulation without phenol red for 5 min at room temperature in the dark. After staining, the solution was replaced with fresh choline-free Medium 199 (without phenol red) containing the same supplements as used during the 4 h incubation. Imaging of live cells was performed using Olympus FV3000 equipped with a 60× oil-immersion objective (UPlanSApo 60×/1.35 Oil, Olympus). FM4-64 and Hoechst 33342 were excited at 561 and 405 nm, respectively, and fluorescence was collected using standard emission filters. All imaging was completed within 40 min after staining.

### Muscle stem cells

C57BL/6 J mice (6–8 weeks, female) were purchased from Japan SLC, Inc. All animal husbandry and experimental procedures were approved by the Animal Care Use and Review Committee of University of Shizuoka. Muscle stem cells were isolated from mouse limb muscles following established protocols^55^. In brief, skeletal muscle tissues (tibialis anterior, quadriceps, extensor digitorum longus, gastrocnemius, plantaris, and soleus) from the limbs were digested with 0.2% collagenase type I to obtain mononuclear cells. The resulting cell suspension was incubated with phycoerythrin (PE)-conjugated anti-mouse Ly-6A/E (Sca-1) (1:200), CD45 (1:200), and CD31 (1:200) antibodies, together with allophycocyanin (APC)-conjugated anti-mouse CD106 (1:100), for 30 min at 4°C. After washing with PBS supplemented with 2% FBS, CD106-positive cells were purified by flow cytometry using an MA900 (Sony). MuSCs were cultured in growth medium consisting of DMEM supplemented with 30% fetal bovine serum, 1% chicken embryo extract, 10 ng/mL basic fibroblast growth factor, and 1% penicillin–streptomycin. Culture dishes were pre-coated with Matrigel. For induction of differentiation, growth medium was removed and washed with PBS for 3 times and replaced with synthetic differentiation medium composed of choline-free DMEM supplemented with 2% B27, 1% penicillin–streptomycin, and choline chloride or choline analogs. The cells were subsequently differentiated for an additional 4 days. For Immunofluorescence imaging, samples were fixed with 4% paraformaldehyde at room temperature for 10 min, permeabilized with 0.5% Triton X-100 for 10 min and then blocked with 1% BSA in PBS for 1 h at room temperature. The samples were incubated with the primary antibody (Myosin 4 Monoclonal Antibody, MF20, 1:500) overnight at 4°C, followed by detection with the secondary antibody (1:500). Nuclei were counterstained with 4′,6-diamidino-2-phenylindole (DAPI) (1:500). Morphology of cells labeled with fluorescent dyes were analyzed by an epifluorescence microscope (Axio-observer Z1).

### Yeast

Using a wild-type yeast strain (YKT38, *MATa lys2-801 ura3-52 his3*Δ-*200 trp1*Δ-*63 leu2*Δ-*1*)^58^, *opi3 cho2* (YKT1963, *MATa cho2*Δ*::HIS3MX6 opi3*Δ*::KanMX6 TRP1 lys2*) was generated by standard yeast genetic manipulation according to previously reported methods^59^. The strain maintained on YPD agar plates was inoculated into choline-free synthetic defined (SD) medium and cultured overnight at 30°C with shaking at 150 rpm. The optical density of the cell suspension was measured at 600 nm (OD₆₀₀), and cells were diluted in SD medium containing the indicated concentrations of choline chloride or choline analogs to an initial OD₆₀₀ of 0.01. Cultures were then incubated at 30°C with shaking at 150 rpm for 48 h. Cell viability was determined using the CellTiter-Glo^®^ 2.0 ATP assay according to the manufacturer’s instructions.

For qPCR analysis, a single colony was inoculated in YPD medium and overnight at 30°C with shaking at 150 rpm to the stationary phase. The cells were diluted in SD medium containing the indicated concentrations of choline chloride or choline analogs to OD₆₀₀ of 0.1 and were grown for additional 24 h. Total RNA was extracted using the Qiagen RNeasy Mini kit. Briefly, cultures were centrifuged at 1,000 × g for 5 min at 4°C. Following removal of the supernatant, lysis buffer and acid-washed glass beads were added to the pellets (at least 5 OD). This mixture was vortexed for 1 min and immediately incubated on ice for 1 min. After repeating the above steps four times, the mixture was centrifuged at maximum speed for 2 min to obtain the total RNA. The remaining procedures were performed according to the manufacturer’s instructions. cDNA was synthesized using the Prime Script Ⅱ First strand cDNA synthesis kit. Quantitative PCR was carried out using a LightCycler^®^ 96 qPCR machine and Powere Up SYBR Green Master Mix. Relative expression levels were calculated by 2^−Δ^ ^Δ Ct^ method with *Alg9* as an internal reference gene. We used these primers^60,61^; *Hac1*_spliced_ forward primer: 5-GCGTAATCCAGAAGCGCAGT-3, *Hac1*_spliced_ reverse primer: 5-GTGATGAAGAAATCATTCAATTCAAATG-3, *Alg9* forward primer: 5-CACGGATAGTGGCTTTGGTGAACAATTAC-3, *Alg9* reverse primer: 5- TATGATTATCTGGCAGCAGGAAAGAACTTGGG-3 The qPCR program included Step 1: 95°C, 20 s; Step 2: 58°C, 20 s; Step 3: 72°C, 30 s, Step 4: 72°C, 5 min; the steps 1–3 were carried out in 40 cycles.

### MD simulation

In this study, we employed 12 bilayer systems, each composed of different xenobiotic phospholipids. Here, we used xenobiotic phospholipids in which the choline group of 1-palmitoyl-2-oleoyl-glycero-3-phosphocholine (POPC) molecules was replaced with a protonated choline analog (Fig. 4h). The force fields for POPC and 1-palmitoyl-2-oleoyl-sn-glycero-3-phosphoethanolamine (POPE) were taken from CHARMM36^62^. For the other xenobiotic phospholipids, the force fields were generated using the CHARMM General Force Field (CGenFF)^63^, and the parameters for regions other than the choline analog part were replaced with the corresponding values from POPC in CHARMM36. The bonded parameters across the junction between the phosphate group and the choline analog were adopted from the CGenFF-generated values. Non-bonded interactions (Lennard-Jones potentials) between atoms from CHARMM36 and CGenFF were generated according to the Lorentz–Berthelot rules^64^. To correct for deviations in the total charge caused by partial substitution of the force field, the charges on the phosphate group were manually adjusted by up to 0.3%. The TIP3P model^65^ was used for water molecules. The POPC and POPE bilayer systems were constructed using the CHARMM-GUI Membrane Builder^66^. The other 11 systems were constructed using a method inspired by the InflateGRO methodology^67^. Xenobiotic phospholipids were initially arranged in a bilayer configuration with sufficiently large intermolecular distances, followed by iterative compression and energy minimization steps. Once an appropriate membrane area was achieved, water molecules were inserted into the non-membrane regions. Each system consisted of 128 lipid molecules and 6,400 water molecules. All systems were equilibrated under conditions of 310.15 K and 0.1 MPa for at least 600 ns, and the final 500 ns were used for analysis. The MD simulation parameters were the same to those generated by the CHARMM-GUI Membrane Builder^66^. The leap-frog algorithm was used for the numerical solution of the equations of motion, and the time step was set to 2.0 fs. The particle mesh Ewald method^68^ with periodic boundary conditions in all directions was used to calculate the long-range Coulomb interactions with a maximum Fourier spacing of 0.12 nm and a fourth-order fitting function. The short-range Coulomb interactions were cut off at 1.2 nm. The van der Waals interactions were smoothly shifted to 0 between 1.0 and 1.2 nm. All of the bonds with hydrogen atoms were constrained using the linear constraint solver^69^. The temperatures of lipid and water were separately kept constant using the velocity rescaling method^70^ with a 1.0-ps coupling constant. The pressure normal and lateral to the bilayer plane were separately maintained using the stochastic cell rescaling method^71^ with a 5.0-ps coupling constant. The centers of mass motion of the membrane and solvent were separately removed every 100 steps. All MD simulations were performed using GROMACS 2024^72^, and all snapshots were rendered using Visual Molecular Dynamics (VMD)^73^.

As characteristic features of the bilayers composed of xenobiotic phospholipids, the following 12 values were calculated. To describe the overall structural and mechanical properties of the bilayer, the area per lipid *a*_0_, the distance *h*_P_ from the bilayer midplane to the phosphorus atom along the bilayer normal, and the area compressibility modulus *K_a_*, were evaluated. The value of *K_a_* was calculated from the bilayer area fluctuations^74^. As properties related to the choline analog part, the PN vector connecting the phosphorus and nitrogen atoms in the lipid headgroup, as well as the radial distribution function (RDF) between the nitrogen atoms in the lipid headgroup and the oxygen atoms of water molecules were analyzed. The mean and standard deviation of the absolute value of the bilayer-normal component of the PN vector were defined as *h*_PN_ and *dh*_PN_, respectively, while those of the absolute value of the bilayer-parallel component were defined as *w*_PN_ and *dw*_PN_. The angle between the PN vector and the bilayer normal vector pointing toward the outside of the bilayer was denoted as *θ*_PN_. For the RDF, the peak position *r*_1_ and peak height *g*(*r*_1_) in the range 0.25–0.35 nm, as well as the peak position *r*_2_ and peak height *g*(*r*_2_) in the range 0.40–0.53 nm, were used as characteristics indicators.

### Statistics and reproducibility

The quantitative data were represented as mean and standard error of the mean. Statistical analyses were performed using one-way ANOVA. Sample sizes were included in figures or legends.

## Appendix

Movie S1. Time-lapse images of **A10**-treated cardiomyocytes.

Cardiomyocytes isolated from rat ventricles were cultured with **A10** (1 mM) in M199 media for 4 hours, and subjected to bright filed imaging (video speed, real time; scale bar, 50 μm).

Movie S2. Time-lapse images of **A13**-treated cardiomyocytes.

Cardiomyocytes isolated from rat ventricles were cultured with **A13** (1 mM) in M199 media for 4 hours, and subjected to bright filed imaging (video speed, real time; scale bar, 50 μm).

Excel file

This contains information of medium formulations (Sheet1), information of reagents and resources (Sheet2), data of MS-based lipid analysis (Sheet3), data of GO term analysis (Sheet4), data of transcriptome analysis (Sheet5), data of interactome analysis (Sheet6), summary data of choline analogs (Sheet7), and individual data of figure items (Sheet8).

**Figure S1.**
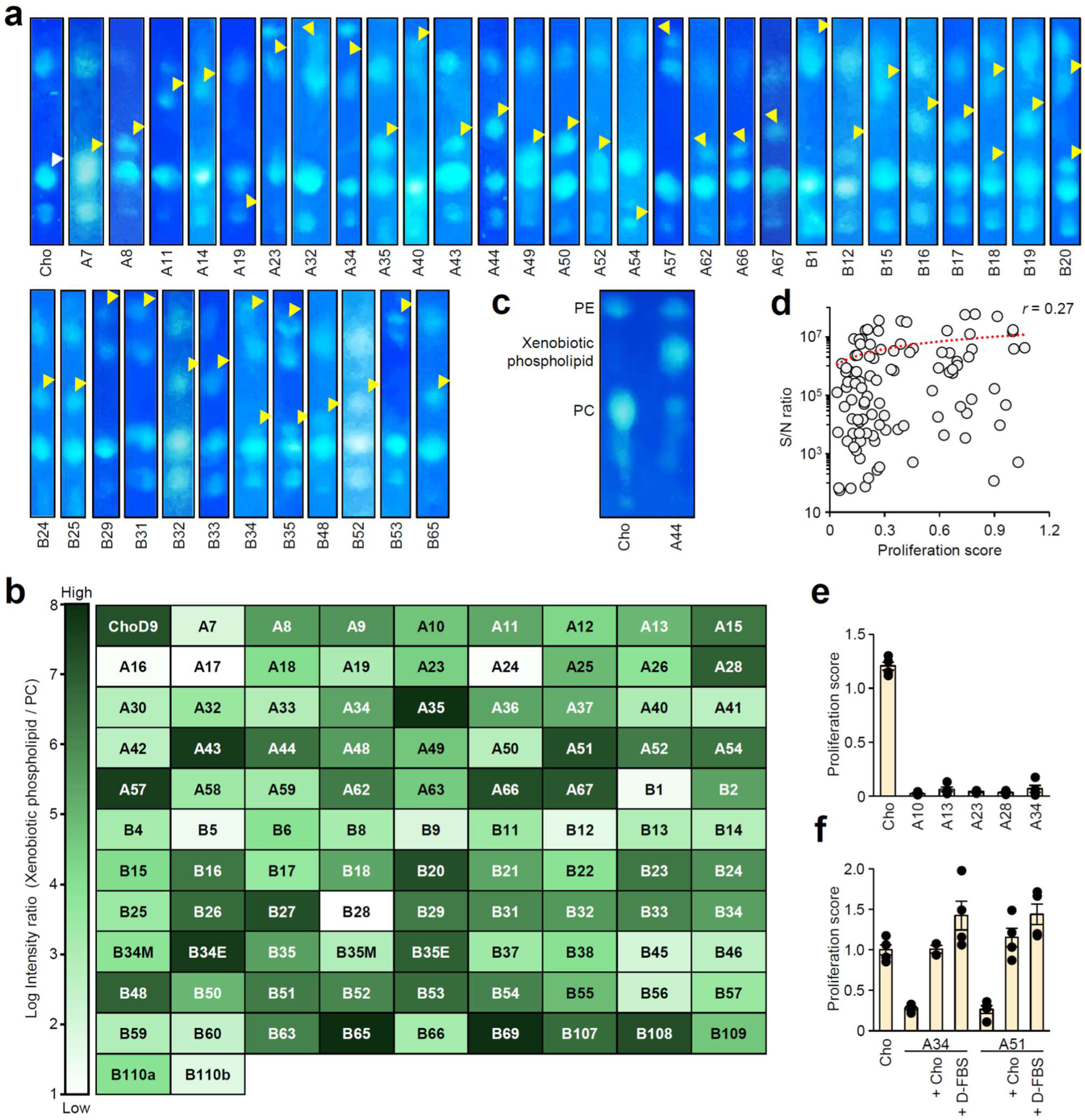
Supporting data for TLC and MS analyses of xenobiotic phospholipids. (**a**) TLC-based detection of xenobiotic phospholipids. K562 cells cultured with analogs (30 μM) in choline-depleted SMv1 for one day were subjected to lipid extraction, TLC separation and primulin staining. TLC images show specific spots (yellow arrowheads) separated from endogenous PC (white arrowhead). (**b**) MS-based detection of xenobiotic phospholipids. K562 cells cultured with analogs (30 μM) in choline-depleted SMv1 for one day were subjected to lipid extraction followed by LC-MS/MS analysis. The ion signals from target phospholipid species (32:1, 32:2, 34:1, 34:2, 36:1, 36:2) were identified by their characteristic ion fragmentation patterns. For each species, the ion intensity of analog-treated cells was compared to that of choline-supplemented control to determine the S/N ratio. The S/N ratios of the top three species in each phospholipid class are averaged and presented in the heatmap table. Detailed m/z parameters and individual data are provided in Excel file. (**c**) Predominant production of xenobiotic phospholipids. K562 cells cultured with **A44** (30 μM) in choline-depleted SMv1 for three days were subjected to TLC analysis. (**d**) Correlation analysis between S/N ratios (Fig. S1b) and proliferation scores (Fig. 2b and 3). The *r* value was determined by linear regression analysis. (**e**) Unchanged non-proliferative effects under a high analog concentration. K562 cells cultured with analogs (500 μM) in choline-depleted SMv1 were analyzed by cell counting (mean ± SEM, n = 4 biological replicates). (**f**) Counteracting non-proliferative effects of analogs by choline and dialyzed serum. K562 cells cultured with analogs (30 μM) and together with choline (500 μM) or dialyzed FBS (D-FBS, 5%) in choline-depleted SMv1 were analyzed by cell counting (mean ± SEM, n = 2–4 biological replicates).

**Figure S2.**
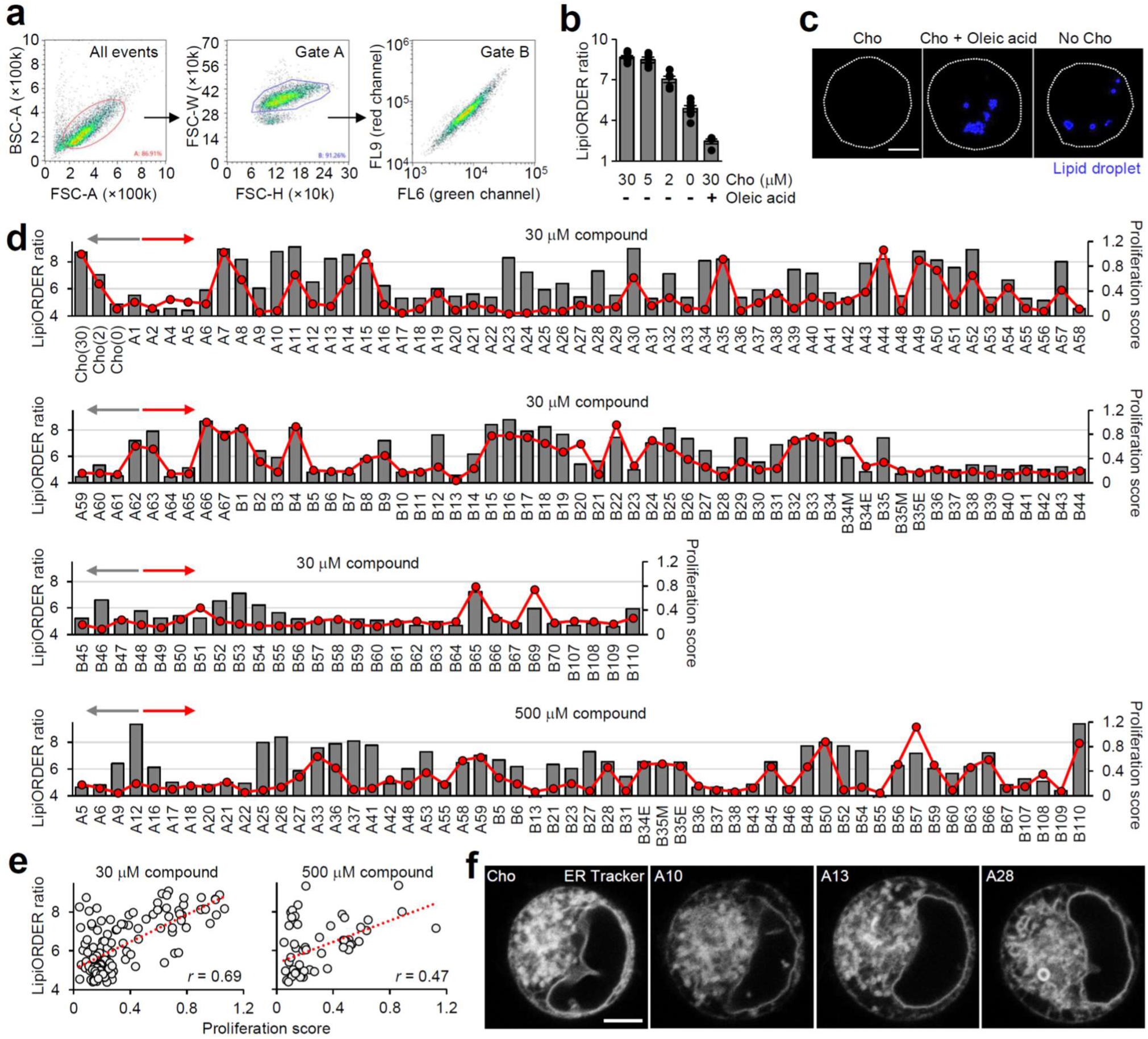
Supporting data for LipiORDER analysis of analog-treated cells. (**a**) Gating strategy for flowcytometric LipiORDER analysis. Live K562 cells were distinguished from cell debris and dead cells by forward scatter (FSC) and back scatter (BSC) (All events). Singlet cells were separated from doublet cells by physical parameters (FSC-H vs FSC-W, Gate A). Fluorescence of LipiORDER excited by a 405-nm laser was measured through FL6 (green) and FL9 (red) channels equipped with band path filters 450/50 and 617/30 nm, respectively (Gate B). (**b**) Decreases in LipiORDER red/green ratio by choline depletion and fatty acid loading. K562 cells cultured with choline (0–30 μM) or oleic acid-albumin complex (100 μM) in SMv1 for one day were stained with LipiORDER and analyzed by flowcytometry to determine the fluorescence intensity ratio (FL9/FL6 in Gate B of Fig. S2a). Bar graphs represent mean ± SEM (n = 4–9 biological replicates). (**c**) Lipid droplet accumulation in choline-depleted cells. K562 cells cultured with choline (0 or 30 μM) or oleic acid-albumin complex (100 μM) in SMv1 for one day were stained with LipiDye II and analyzed by live-cell confocal microscopy. White lines represent the periphery of cells. (**d**) LipiORDER ratios and proliferation scores in analog-treated cells. K562 cells cultured with choline (0, 2 or 30 μM) or analogs (30 or 500 μM) in choline-depleted SMv1 for one day were subjected to flowcytometric LipiORDER analysis as described in Fig. S2a and S2b. Bar graphs show the average LipiORDER ratios (n = 2–9 biological replicates) and overlayed dotted lines show the average proliferation scores (n = 3–33 biological replicates). (**e**) Correlation analysis between LipiORDER ratios and proliferation scores (data from Fig. S2d). The *r* values were determined by linear regression analysis. (**f**) ER structures in analog-treated cells. K562 cells cultured with analogs (500 μM) for two days were stained with ER-tracker and analyzed by live-cell confocal microscopy. Scale bars: **c**, **f** 5 µm.

**Figure S3.**
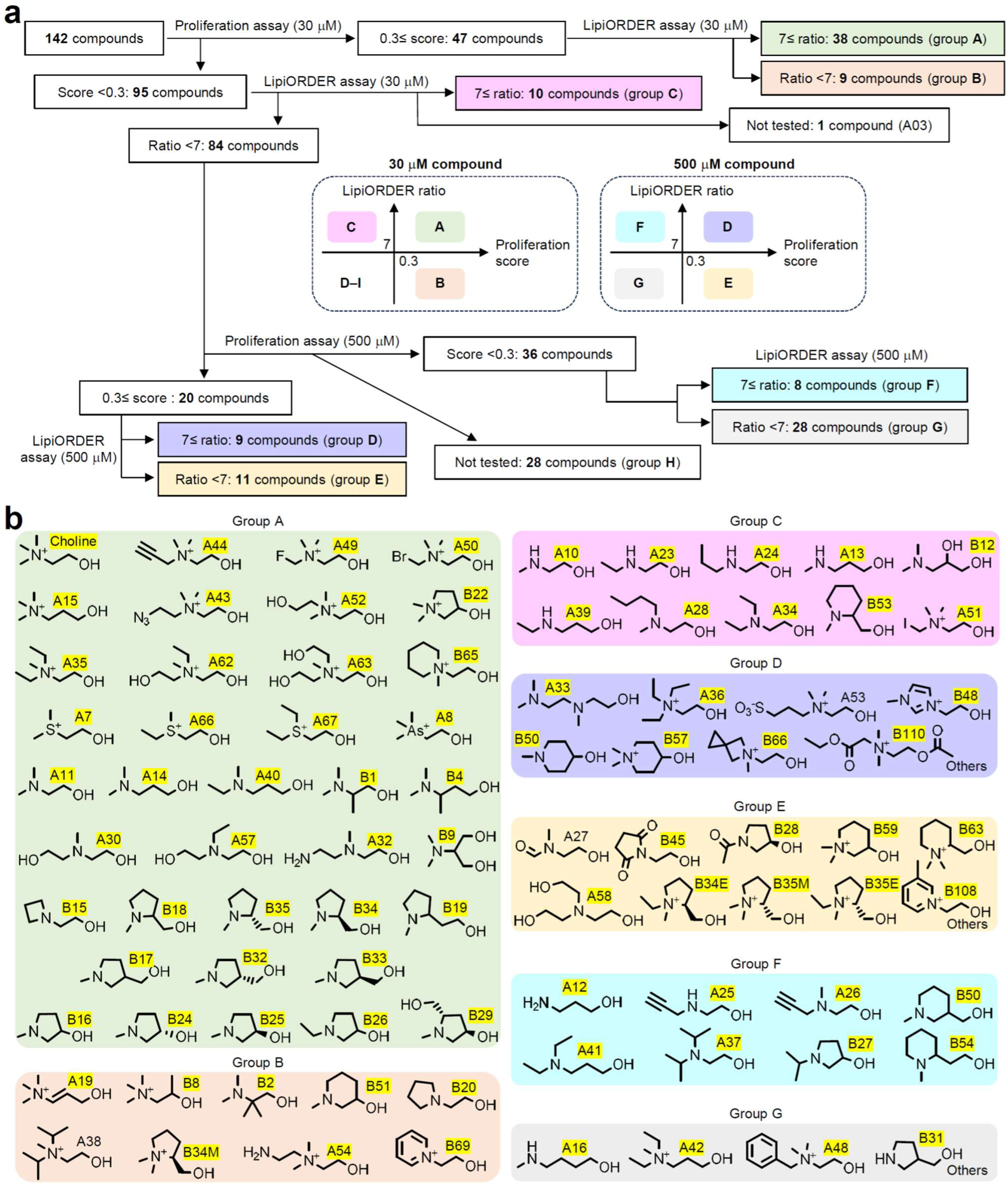
Supporting data for classification of choline analogs by proliferation scores and LipiORDER ratios. (**a**) Flowchart showing steps for classification of 142 compounds based on the results of cell proliferation and LipiORDER assays (Fig. 2b, 3, S2d). (**b**) The compounds are classified according to **a** (groups highlighted in different background colors). Compound ID is highlighted in yellow when its xenobiotic phospholipids were detected by MS (S/N >10) or TLC. Detailed data are provided in Excel file.

**Figure S4.**
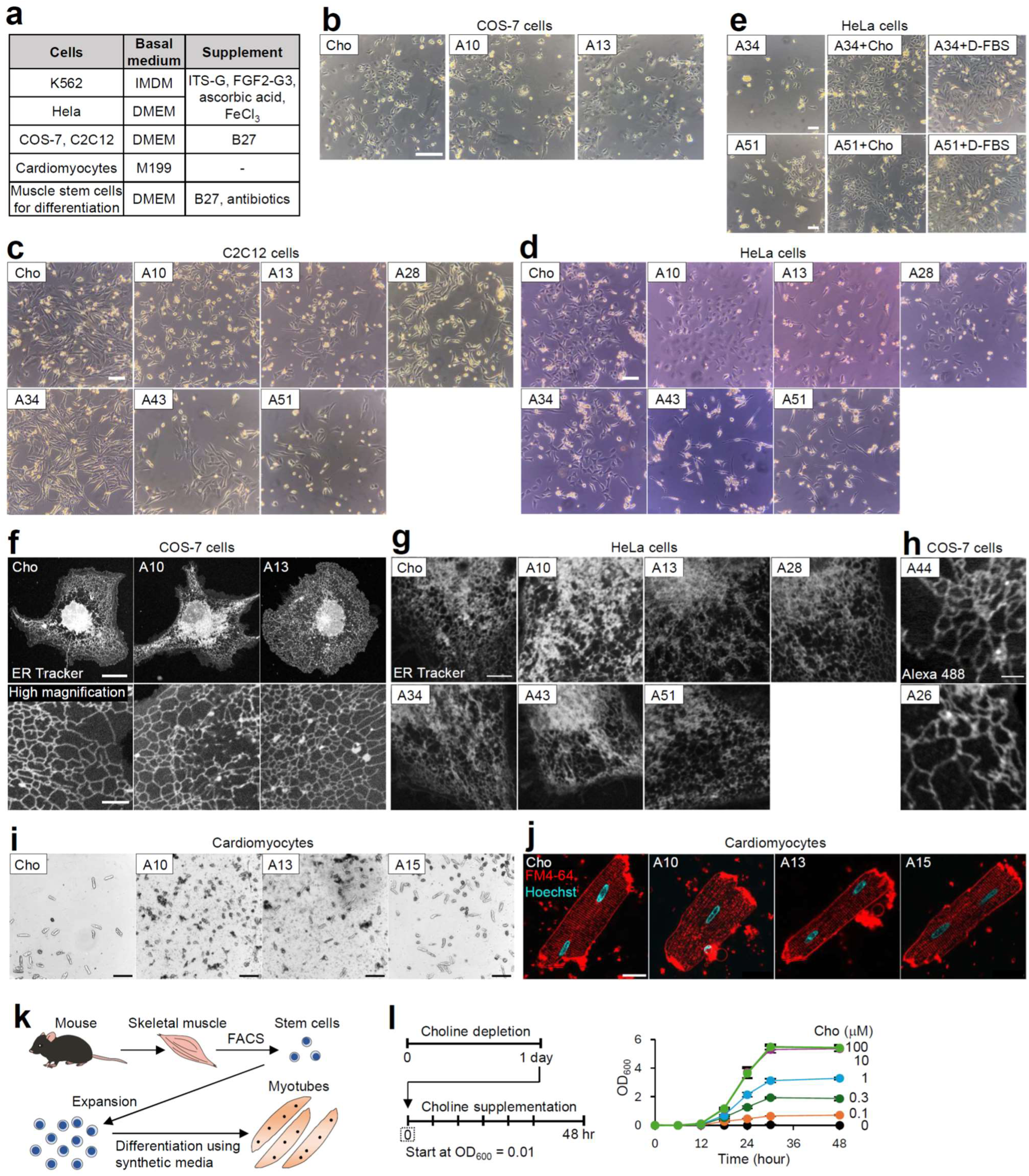
Supporting data for culture of adherent cell lines, primary cells and yeast cells in synthetic media. (**a**) Synthetic media used for mammalian cells in this study (detailed composition in Excel file). (**b–d**) Bright field images of adherent cell lines cultured with choline and analogs (500 μM) in corresponding synthetic media (Fig. S4a) for four days (**b**, COS-7; **d**, HeLa) or two days (**c**, C2C12). (**e**) Counteracting non-proliferative effects of analogs by supplementation with choline and dialyzed serum. HeLa cells cultured with analogs (30 μM) and together with choline (500 μM) or D-FBS (10%) in choline-depleted SMv1 were analyzed by bright-field microscopy. (**f, g**) ER structures in analog-treated cells. Adherent cell lines cultured with choline and analogs (500 μM) in synthetic media for three or four days were stained with ER-tracker and analyzed by live-cell confocal microscopy (**f**, COS-7; **g**, HeLa). (**h**) Alkyne-containing xenobiotic phospholipids resident in ER membranes. COS-7 cells cultured with analogs (30 μM) in synthetic media for one day were labeled with Alexa Fluor 488 azide through copper-catalyzed azide–alkyne cycloaddition, followed by confocal microscopy. (**i**) Bright field images of rat cardiomyocytes cultured with choline and analogs (1 mM) in M199 medium for 4 hours. (**j**) Confocal imaging of T-tubule structure in analog-treated cardiomyocytes. Cardiomyocytes were cultured with choline and analogs (1 mM) in M199 medium for 4 hours, and stained with FM4-64 and Hoechst 33342 for live-cell confocal microscopy. (**k**) Workflow for inducing myogenic differentiation of muscle stem cells with synthetic media. (**l**) Choline-dependent growth of *cho2Δopi3Δ* cells in SD medium. The yeast cells pre-cultured in choline-depleted SD medium for one day were cultured in choline-supplemented SD medium for monitoring of growth curves by OD_600_ measurements. Data are mean ± SEM (n = 3 biological replicates). Scale bars: **b–e**, 100 µm; **f**, 20 µm; **f** (high magnification), 5 µm; **g**, 5 µm; **h**, 2 µm; **i**, 500 µm; **j**, 20 µm.

**Figure S5.**
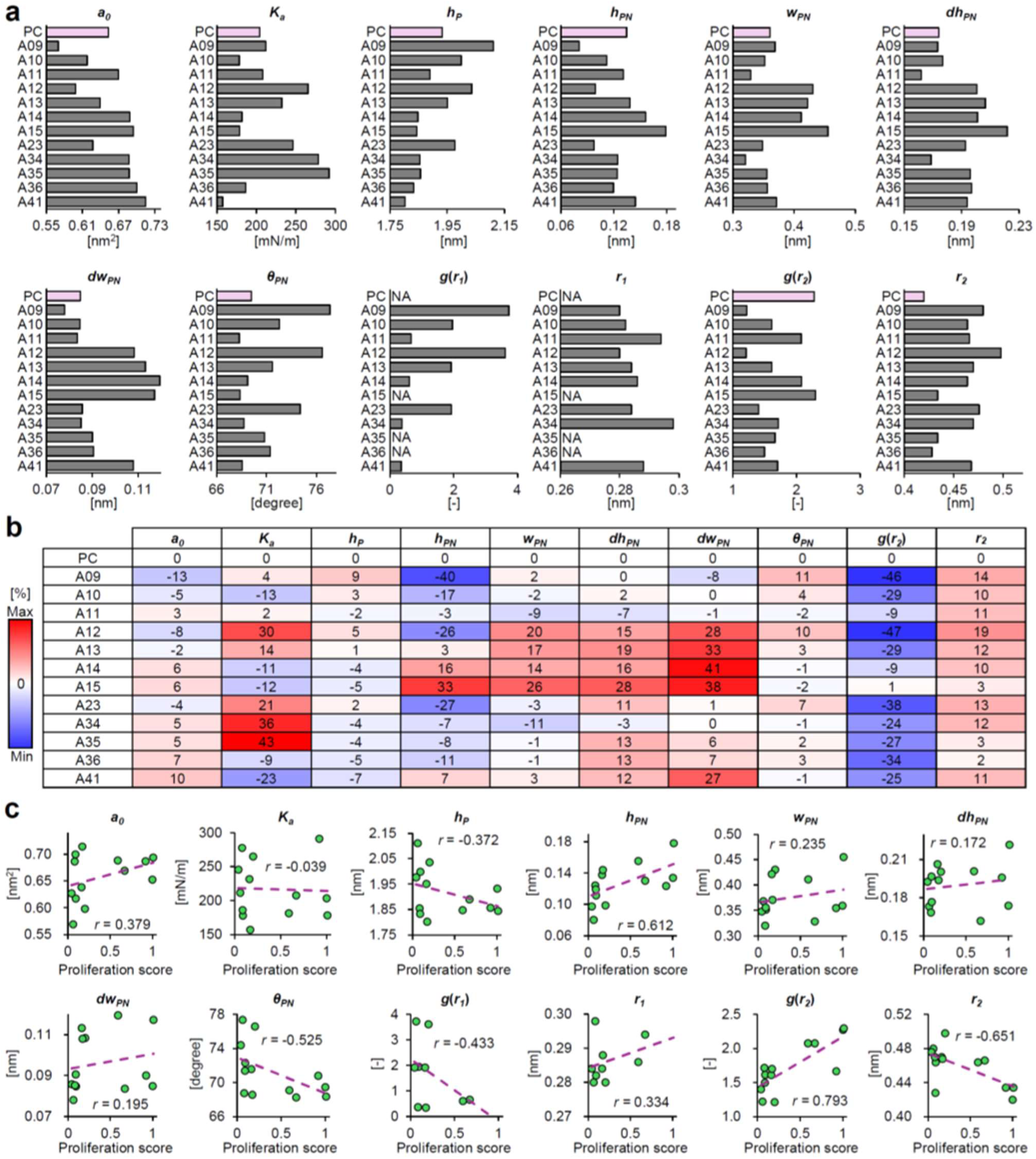
Supporting data for MD simulation of phospholipid bilayer properties. (**a**) Bar graphs showing results of calculating bilayer properties indicated in Fig. 4f. (**b**) Heatmap table showing the percentage change in bilayer properties (Fig. S5a) of headgroup-modified phospholipids compared with those of PC. (**c**) Dot plots representing the bilayer property (Fig. S5a) and proliferation score (Fig. 2b and 3) of each phospholipid. The *r* values were calculated from linear regression models.

**Figure S6.**
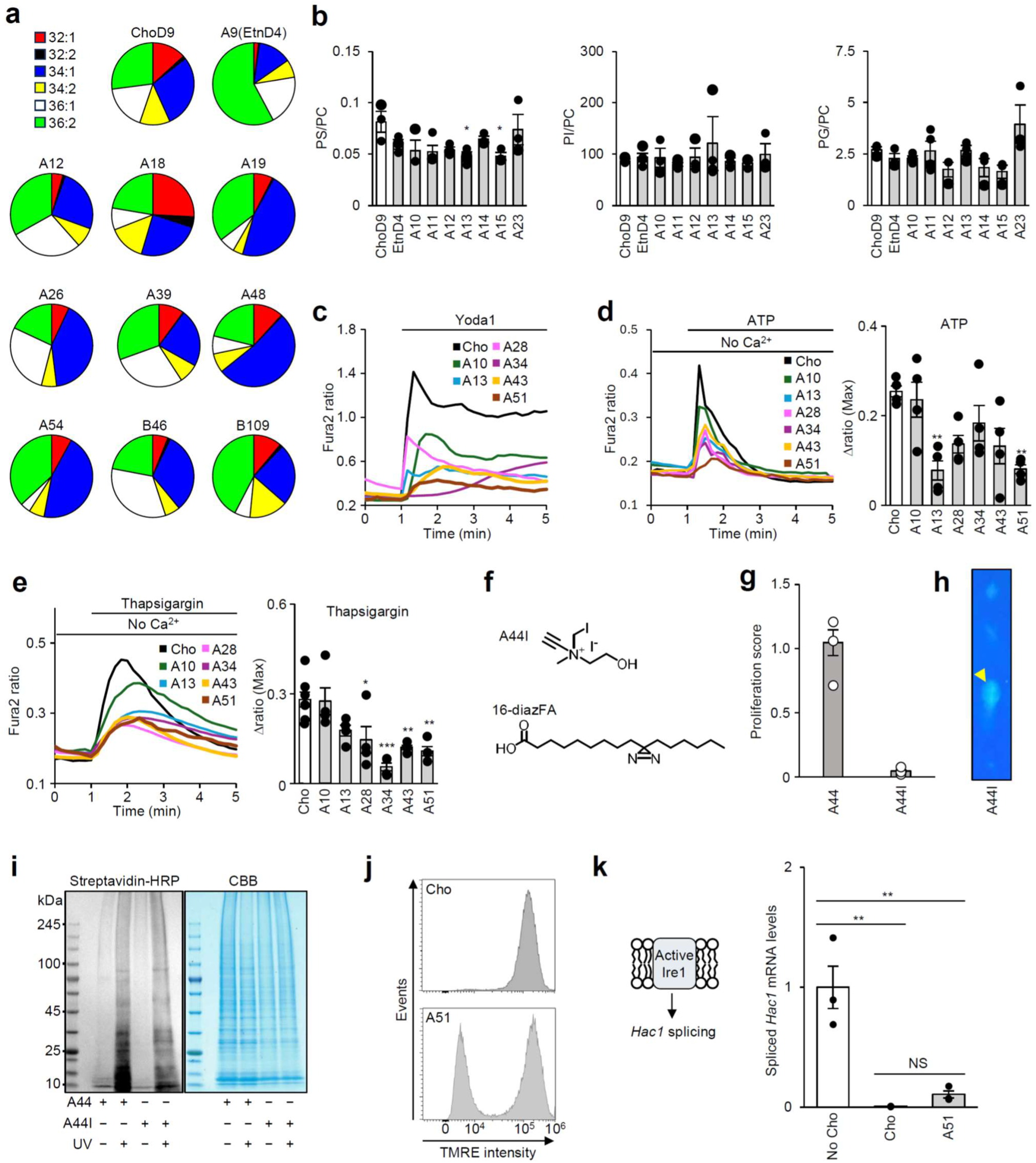
Supporting data for differential biological effects of xenobiotic phospholipids. (**a**) Pie charts showing the acyl chain composition of xenobiotic phospholipids. MS analysis was conducted as described in Fig. S1b (n = 2 or 3 biological replicates, average values shown). Detailed data are provided in Excel file. (**b**) Endogenous lipid composition in analog-treated cells. MS analysis was conducted as described in Fig. S1b, but the targets are changed to endogenous phospholipids (n = 3 or 4 biological replicates). (**c**) Inhibition of PIEZO1-mediated Ca^2+^ influx in analog-treated cells. C2C12 cells cultured with analogs (500 μM) for two days in synthetic media (Fig. S4a) were subjected to fura2 ratiometric imaging with Yoda1. (**d**) Inhibition of ATP-induced Ca^2+^ release in analog-treated cells. HeLa cells cultured with analogs (500 μM) for four days in synthetic media (Fig. S4a) were subjected to fura2 ratiometric imaging with ATP. ATP-induced Ca^2+^ release was quantified as the maximal increment of F_340_/F_380_ (Δ ratio) (n = 4 biological replicates). (**e**) Inhibition of thapsigargin-induced Ca^2+^ release in analog-treated cells. Fura2 imaging was performed as described in Fig. S6d, but thapsigargin was used as a substitute for ATP (n = 4–7 biological replicates). (**f**) Structures of **A44I** and 16-diazFA. (**g**) Non-proliferative activity of **A44I**. K562 cells cultured with **A44** or **A44I** (500 μM) in choline-depleted SMv1 were analyzed by cell counting (n = 3 biological replicates). (**h**) TLC analysis of **A44I**-incorporatred xenobiotic phospholipids. K562 cells cultured with **A44I** (500 μM) in choline-depleted SMv1 were analyzed by TLC (specific spot indicated by yellow arrowhead). (**i**) Photo-crosslinking of xenobiotic phospholipids and interacting proteins. K562 cells cultured with **A44** or **A44I** (500 μM) together with 16-diazFA (100 μM) in choline-depleted SMv1 for one day were subjected to UV irradiation, cell lysis, click labeling with biotin, SDS-PAGE separation, and western blotting using HRP-conjugated streptavidin (left) or CBB staining (right). (**j**) Mitochondrial depolarization in **A51**-treated cells. K562 cells cultured with choline or **A51** (500 μM) in choline-depleted SMv1 for three days were stained with TMRE, followed by flow cytometry. (**k**) Real-time PCR analysis of *cho2Δopi3Δ* cells cultured with choline or analogs (0 or 100 μM) in SD medium. Spliced *Hac1* mRNA levels are normalized to the *Alg9* reference gene (n = 3 biological replicates). Bar graphs represent mean ± SEM. \**p* < 0.05, \*\**p* < 0.01, \*\*\**p* < 0.001 (one-way ANOVA).

## Notes

### Competing Interest Statement

The authors have declared no competing interest.

## References

1. Van Meer, G., Voelker, D. R. & Feigenson, G. W. Membrane lipids: Where they are and how they behave. Nat. Rev. Mol. Cell Biol. 9, 112–124 (2008).

2. Harayama, T. & Riezman, H. Understanding the diversity of membrane lipid composition. Nat. Rev. Mol. Cell Biol. 19, 281–296 (2018).

3. Winnikoff, J. R. et al. Homeocurvature adaptation of phospholipids to pressure in deep-sea invertebrates. Science 384, 1482–1488 (2024).

4. Yeung, T. et al. Membrane phosphatidylserine regulates surface charge and protein localization. Science 319, 210–213 (2008).

5. Laganowsky, A. et al. Membrane proteins bind lipids selectively to modulate their structure and function. Nature 510, 172–175 (2014).

6. Crotta Asis, A., Asaro, A. & D’Angelo, G. Single cell lipid biology. Trends Cell Biol. 35, 651–666 (2025).

7. Kenny, T. C., Scharenberg, S., Abu-Remaileh, M. & Birsoy, K. Cellular and organismal function of choline metabolism. Nat. Metab. 7, 35–52 (2025).

8. Sohlenkamp, C. & Geiger, O. Bacterial membrane lipids: Diversity in structures and pathways. FEMS Microbiol. Rev. 40, 133–159 (2015).

9. Bao, X. et al. Shortening of membrane lipid acyl chains compensates for phosphatidylcholine deficiency in choline-auxotroph yeast. EMBO J. 40, e107966 (2021).

10. Tavasoli, M., Lahire, S., Reid, T., Brodovsky, M. & McMaster, C. R. Genetic diseases of the Kennedy pathways for membrane synthesis. J. Biol. Chem. 295, 17877–17886 (2020).

11. Chen, X. W. et al. Time for lipid cell biology. Nat. Cell Biol. 27, 169–174 (2025).

12. Chen, P. H. B., Li, X. L. & Baskin, J. M. Synthetic Lipid Biology. Chem. Rev. 125, 2502–2560 (2025).

13. Li, G. et al. Efficient replacement of plasma membrane outer leaflet phospholipids and sphingolipids in cells with exogenous lipids. Proc. Natl. Acad. Sci. U. S. A. 113, 14025–14030 (2016).

14. Tei, R., Bagde, S. R., Fromme, J. C. & Baskin, J. M. Activity-based directed evolution of a membrane editor in mammalian cells. Nat. Chem. 15, 1030–1039 (2023).

15. Tamura, T. et al. Organelle membrane-specific chemical labeling and dynamic imaging in living cells. Nat. Chem. Biol. 16, 1361–1367 (2020).

16. Tsuchiya, M., Tachibana, N., Nagao, K., Tamura, T. & Hamachi, I. Organelle-selective click labeling coupled with flow cytometry allows pooled CRISPR screening of genes involved in phosphatidylcholine metabolism. Cell Metab. 35, 1072–1083 (2023).

17. Cantor, J. R. et al. Physiologic Medium Rewires Cellular Metabolism and Reveals Uric Acid as an Endogenous Inhibitor of UMP Synthase. Cell 169, 258–272 (2017).

18. Kenny, T. C. et al. Integrative genetic analysis identifies FLVCR1 as a plasma-membrane choline transporter in mammals. Cell Metab. 35, 1057–1071 (2023).

19. Scharenberg, S. G. et al. An SPNS1-dependent lysosomal lipid transport pathway that enables cell survival under choline limitation. Sci. Adv. 9, eadf8966 (2023).

20. Chen, G. et al. Chemically defined conditions for human iPSC derivation and culture. Nat. Methods 8, 424–429 (2011).

21. Chen, S. et al. Role of Gpcpd1 in intestinal alpha-glycerophosphocholine metabolism and trimethylamine N-oxide production. J. Biol. Chem. 300, 107965 (2024).

22. Malito, E., Sekulic, N., Too, W. C. S., Konrad, M. & Lavie, A. Elucidation of Human Choline Kinase Crystal Structures in Complex with the Products ADP or Phosphocholine. J. Mol. Biol. 364, 136–151 (2006).

23. Son, Y., Kenny, T. C., Khan, A., Birsoy, K. & Hite, R. K. Structural basis of lipid head group entry to the Kennedy pathway by FLVCR1. Nature 629, 710–716 (2024).

24. Jao, C. Y., Roth, M., Welti, R. & Salic, A. Metabolic labeling and direct imaging of choline phospholipids in vivo. Proc. Natl. Acad. Sci. U. S. A. 106, 15332–15337 (2009).

25. Jao, C. Y., Roth, M., Welti, R. & Salic, A. Biosynthetic labeling and two-color imaging of phospholipids in cells. ChemBioChem 16, 472–476 (2015).

26. Jiang, G., et al. Chemical Approaches to Optimize the Properties of Organic Fluorophores for Imaging and Sensing. Angew. Chemie - Int. Ed. 63, e202315217 (2024).

27. Wang, D., et al. Global Mapping of Protein–Lipid Interactions by Using Modified Choline-Containing Phospholipids Metabolically Synthesized in Live Cells. Angew. Chemie - Int. Ed. 56, 5829–5833 (2017).

28. Valanciunaite, J. et al. Polarity Mapping of Cells and Embryos by Improved Fluorescent Solvatochromic Pyrene Probe. Anal. Chem. 92, 6512–6520 (2020).

29. Luo, M. Chemical and Biochemical Perspectives of Protein Lysine Methylation. Chem. Rev. 118, 6656–6705 (2018).

30. Xiao, B. Mechanisms of mechanotransduction and physiological roles of PIEZO channels. Nat. Rev. Mol. Cell Biol. 25, 886–903 (2024).

31. Prakriya, M. & Lewis, R. S. Store-operated calcium channels. Physiol. Rev. 95, 1383–1436 (2015).

32. Pakos-Zebrucka, K. et al. The integrated stress response. EMBO Rep. 17, 1374–1395 (2016).

33. Tang, D., Kang, R., Berghe, T. Vanden, Vandenabeele, P. & Kroemer, G. The molecular machinery of regulated cell death. Cell Res. 29, 347–364 (2019).

34. Wilcken, R., Zimmermann, M. O., Lange, A., Joerger, A. C. & Boeckler, F. M. Principles and applications of halogen bonding in medicinal chemistry and chemical biology. J. Med. Chem. 56, 1363–1388 (2013).

35. Nah, J. et al. Microprotein SMIM26 drives oxidative metabolism via serine-responsive mitochondrial translation. Mol. Cell 85, 2759–2775 (2025).

36. Hayashi, T. et al. Higd1a is a positive regulator of cytochrome c oxidase. Proc. Natl. Acad. Sci. U. S. A. 112, 1553–1558 (2015).

37. Alahmad, A., et al. Bi-allelic pathogenic variants in NDUFC2 cause early-onset Leigh syndrome and stalled biogenesis of complex I. EMBO Mol. Med. 12, e12619 (2020).

38. Zhang, S. et al. Mitochondrial peptide BRAWNIN is essential for vertebrate respiratory complex III assembly. Nat. Commun. 11, 1312 (2020).

39. Ko, S. H., Cho, B. L. & Shin, D. Microproteins in Metabolic Biology: Emerging Functions and Potential Roles as Nutrient-Linked Biomarkers. Int. J. Mol. Sci. 26, 11883 (2025).

40. Decker, S. T. & Funai, K. Mitochondrial membrane lipids in the regulation of bioenergetic flux. Cell Metab. 36, 1963–1978 (2024).

41. von Maydell, D. et al. ABCA7 variants impact phosphatidylcholine and mitochondria in neurons. Nature 647, 462–471 (2025).

42. Hong, T. T. & Shaw, R. M. Cardiac t-tubule microanatomy and function. Physiol. Rev. 97, 227–252 (2017).

43. Sun, D. en et al. Click-ExM enables expansion microscopy for all biomolecules. Nat. Methods 18, 107–113 (2021).

44. Cardoso, A. C. et al. Mitochondrial substrate utilization regulates cardiomyocyte cell-cycle progression. Nat. Metab. 2, 167–178 (2020).

45. Relaix, F. et al. Perspectives on skeletal muscle stem cells. Nat. Commun. 12, 692 (2021).

46. Belardi, B., Son, S., Felce, J. H., Dustin, M. L. & Fletcher, D. A. Cell–cell interfaces as specialized compartments directing cell function. Nat. Rev. Mol. Cell Biol. 21, 750–764 (2020).

47. Celik, C., Lee, S. Y. T., Yap, W. S. & Thibault, G. Endoplasmic reticulum stress and lipids in health and diseases. Prog. Lipid Res. 89, 101198 (2023).

48. Bock, C. et al. High-content CRISPR screening. Nat. Rev. Methods Prim. 2, 8 (2022).

49. Timmann, S., Feng, Z. & Alcarazo, M. Recent Applications of Sulfonium Salts in Synthesis and Catalysis. Chem. - A Eur. J. 30, e202402768 (2024).

50. Sowmiah, S., Esperança, J. M. S. S., Rebelo, L. P. N. & Afonso, C. A. M. Pyridinium salts: From synthesis to reactivity and applications. Org. Chem. Front. 5, 453–493 (2018).

51. Chiu, D. C. & Baskin, J. M. Organelle-Selective Membrane Labeling through Phospholipase D-Mediated Transphosphatidylation. JACS Au 2, 2703–2713 (2022).

52. Tsuchiya, M., Tachibana, N. & Hamachi, I. Post-click labeling enables highly accurate single cell analyses of glucose uptake ex vivo and in vivo. *Commun*. Biol. 7, 459 (2024).

53. Sato, T. et al. LPGAT1/LPLAT7 regulates acyl chain profiles at the sn-1 position of phospholipids in murine skeletal muscles. J. Biol. Chem. 299, 104848 (2023).

54. Tsuchiya, M. et al. Cell surface flip-flop of phosphatidylserine is critical for PIEZO1-mediated myotube formation. Nat. Commun. 9, 2049 (2018).

55. Hirano, K. et al. Mg^2+^ influx mediated by TRPM7 triggers the initiation of muscle stem cell activation. Sci. Adv. 11, eadu0601 (2025).

56. Konno, R. et al. Universal Pretreatment Development for Low-input Proteomics Using Lauryl Maltose Neopentyl Glycol. Mol. Cell. Proteomics 23, 100745 (2024).

57. Ogura, Y., Ito, H., Sugita, S., Nakamura, M. & Ujihara, Y. Decrease in Ca^2+^ Concentration in Quail Cardiomyocytes Is Faster than That in Rat Cardiomyocytes. Processes 10, 508 (2022).

58. Misu, K. et al. Cdc50p, a conserved endosomal membrane protein, controls polarized growth in Saccharomyces cerevisiae. Mol. Biol. Cell 14, 730–747 (2003).

59. Guide to yeast genetics and molecular biology. Methods in Enzymology vol. 194 (1991).

60. Halbleib, K. et al. Activation of the Unfolded Protein Response by Lipid Bilayer Stress. Mol. Cell 67, 673–684 (2017).

61. Teste, M. A., Duquenne, M., François, J. M. & Parrou, J. L. Validation of reference genes for quantitative expression analysis by real-time RT-PCR in Saccharomyces cerevisiae. BMC Mol. Biol. 10, 99 (2009).

62. Klauda, J. B. et al. Update of the CHARMM All-Atom Additive Force Field for Lipids: Validation on Six Lipid Types. J. Phys. Chem. B 114, 7830–7843 (2010).

63. Vanommeslaeghe, K. et al. CHARMM general force field: A force field for drug-like molecules compatible with the CHARMM all-atom additive biological force fields. J. Comput. Chem. 31, 671–690 (2010).

64. Lorentz, H. A. Ueber die Anwendung des Satzes vom Virial in der kinetischen Theorie der Gase. Ann. Phys. 248, 127–136 (1881).

65. Jorgensen, W. L., Chandrasekhar, J., Madura, J. D., Impey, R. W. & Klein, M. L. Comparison of simple potential functions for simulating liquid water. J. Chem. Phys. 79, 926–935 (1983).

66. Wu, E. L. et al. CHARMM-GUI membrane builder toward realistic biological membrane simulations. J. Comput. Chem. 35, 1997–2004 (2014).

67. Kandt, C., Ash, W. L. & Peter Tieleman, D. Setting up and running molecular dynamics simulations of membrane proteins. Methods 41, 475–488 (2007).

68. Essmann, U. et al. A smooth particle mesh Ewald method. J. Chem. Phys. 103, 8577–8593 (1995).

69. Hess, B., Bekker, H., Berendsen, H. J. C. & Fraaije, J. G. E. M. LINCS: A Linear Constraint Solver for molecular simulations. J. Comput. Chem. 18, 1463–1472 (1997).

70. Bussi, G., Donadio, D. & Parrinello, M. Canonical sampling through velocity rescaling. J. Chem. Phys. 126, 014101 (2007).

71. Bernetti, M. & Bussi, G. Pressure control using stochastic cell rescaling. J. Chem. Phys. 153, 114107 (2020).

72. Berendsen, H. J. C., van der Spoel, D. & van Drunen, R. GROMACS: A message-passing parallel molecular dynamics implementation. Comput. Phys. Commun. 91, 43–56 (1995).

73. Humphrey, W., Dalke, A. & Schulten, K. VMD: Visual molecular dynamics. J. Mol. Graph. 14, 33–38 (1996).

74. Doktorova, M., LeVine, M. V., Khelashvili, G. & Weinstein, H. A New Computational Method for Membrane Compressibility: Bilayer Mechanical Thickness Revisited. Biophys. J. 116, 487–502 (2019).

